# Validation and performance of a quantitative IgG assay for the screening of SARS-CoV-2 antibodies

**DOI:** 10.1101/2020.06.11.146332

**Authors:** Ana M. Espino, Petraleigh Pantoja, Carlos A. Sariol

## Abstract

The current COVID-19 epidemic imposed an unpreceded challenge to the scientific community in terms of treatment, epidemiology, diagnosis, social interaction, fiscal policies and many other areas. The development of accurate and reliable diagnostic tools (high specificity and sensitivity) is crucial in the current period, the near future and in the long term. These assays should provide guidance to identify immune presumptive protected persons, potential plasma, and/or B cell donors and vaccine development among others. Also, such assays will be contributory in supporting prospective and retrospective studies to identify the prevalence and incidence of COVID-19 and to characterize the dynamics of the immune response. As of today, only thirteen serological assays have received the Emergency Use Authorization (EUA) by the U.S. Federal Drug Administration (FDA). In this work we describe the development and validation of a quantitative IgG enzyme-linked immunoassay (ELISA) using the recombinant SARS-CoV-2 Spike Protein S1 domain, containing the receptor-binding domain (RBD), showing 98% sensitivity, 98.9% specificity and positive and negative predictive values of 100% and 99.2%, respectively. The assay showed to be useful to test for SARS-CoV-2 IgG antibodies in plasma samples from COVID-19-recovered subjects as potential donors for plasmapheresis. This assay is currently under review by the Federal Drug Administration for an Emergency Use Authorization request (Submission Number EUA201115).

## Introduction

The current severe acute respiratory syndrome coronavirus 2 (SARS CoV-2) pandemic and the resulting unprecedented outbreak of coronavirus disease 2019 (COVID-19) have shifted the paradigm for viral research, epidemiology and diagnostic. Both molecular and serological methods have been developed at an extraordinary speed. As of April 2, 2020 only four months after the virus was detected for first time in Wuhan region, 28 companies obtained Emergency Use Authorization (EUA) approvals from US Federal Drug Administration (FDA) for their commercial Reverse Transcription-Polymerase Chain Reaction (RT-PCR) diagnostics. Those assays are intended to detect the virus during the acute phase of the infection, providing no information regarding the immunological status of these patients. By the same time, from the more than 25 rapid serological tests available only one had the EUA granted. These rapid tests are relatively simple to perform and interpret and therefore require limited test operator training. The main drawback of these rapid tests is that the specificity and particularly the sensitivity are lower than the standard Enzyme-linked Immunosorbent Assays (ELISA). As of June 1, 2020 FDA had received more than 198 notifications from manufacturers confirming they have validated and intend to distribute their tests in the market. However only 13 of those tests have indeed the EUA from FDA. Moreover, in May 2020, FDA removed 28 SARS-CoV-2 serological tests from the notification list of tests offered during the COVID-10 emergency for not having an EUA request. Choosing an appropriate test to screen for the presence of humoral immune response to SARS-CoV-2 is critical. Such serologic tests are expected to play a key role in the fight against COVID-19 by helping to identify individuals who had developed an adaptive immune response and may be at lower risk of infection. Also, validated serological tests are needed to confirm which subjects, being confirmed positive for COVID-19, truly developed a substantial humoral immune response and may be considered as plasma donors (1). Different antigens have been used to detect antibodies against another novel coronavirus such as SARS-CoV and MERS-CoV (2–5). From these previous works it can be concluded that spike-derived (S) antigens are more sensitive, specific and accurate than nucleocapsid protein-derived (NP) antigens. Also results from assays using S antigens correlated much better with the neutralizing titers than those using NP antigens (6). A recent work showed the usefulness for the Receptor Binding Domain (RBD) and the full Spike protein to detect SARS-CoV-2 specific antibodies (7) and their correlation with neutralizing antibodies nAb (8). For these reasons we choose to use a recombinant SARS-CoV-2 Spike Protein, S1 domain containing the RBD.

With this work we described the validation of a quantitative ELISA, CovIgG-Assay (https://prsciencetrust.org/the-covigg-assay-kit/), showing a very low background and lack of cross reactivity with other respiratory and non-respiratory pathogens in more than 132 samples collected before June 2019. Also the correlation with three serological tests available in the market is described. Finally, we confirm the usefulness of the assay detecting anti-SARS-CoV-2 antibodies in plasma samples from potential plasma donors. CovIgG-Assay is a useful tool to characterize, quantify and to study the dynamics of the humoral immune response to SARS-CoV-2.

## Materials and Methods

### Study Design

The study population included a total of 181 samples. Forty-nine (49) samples were from individuals with symptomatic infection and positive diagnosis for SARS-CoV-2. Forty-eight (48) were confirmed by RT-PCR tests EUA authorized and one (1) diagnosed by COVID-19 ELISA IgG Antibody Test – Mount Sinai, also EUA authorized. De-identified serum or plasma specimens were obtained from local clinical laboratories and Blood Banks and no personal identifiers were retained. The other 132 de-identified samples had been taken previously to 2019 and belonged to the Virology or the Immunology UPR-RCM serum bank. From these samples, 78 had no previous history of viral, allergic or bacterial infections according to our cross-reactivity panel. Nine (9) were previously diagnosed with Zika, three (3) with Dengue, thirteen (13) with history of respiratory allergies and one (1) with Influenza H1N1. We also included a cross reactivity panel with 28 samples kindly donated by the Centers for Disease Control and Prevention (CDC) Dengue Branch, San Juan, PR. These samples included six (6) positives for Respiratory Syncytial Virus (RSV)-IgM, twelve (12) RT-PCR positive for Influenza A or B, five (5) Zika-IgM positive and five (5) positive for Dengue-IgM. This cross-reactive panel was selected according to the most common viral and respiratory infections affecting our population. Additionally, we tested nine (9) samples from individuals that resulted positive for Mycoplasma-IgM and three (3) positives for Chikungunya, which were collected during COVID-19 pandemic. Although these 12 samples were included in the cross-reactive study they were excluded from the statistical analysis to establish the cut-point and diagnostic specificity/sensitivity of CovIgG-Assay. All samples were stored at −80°C until use.

For comparison with two others serological tests (CoronaCheck and Abbott Architect) holding an EUA, we used a set of nine (9) samples assumed to be positive for IgG and IgM and eighteen (18) assumed to be IgG positive for SARS-CoV-2 antibodies. Those samples were also received de-identified from local laboratories.

### CovIgG-Assay

CovIgG-Assay is an indirect ELISA for quantitative determination of human IgG antibody class, which was optimized by checkerboard titration. Disposable high bind flat-bottomed polystyrene 96-wells microtiter plates (Costar, Corning MA No. 3361) were coated overnight at 4°C with 2μg/ml of recombinant SARS-CoV-2 S1-RBD protein (GenScript No. Z03483-1) in carbonate-bicarbonate buffer (Sigma Aldrich No. 08058). Plates were washed 3 times with phosphate buffered saline (PBS) containing 0.05% Tween-20 (PBST) and blocked for 30 min at 37°C with 250μl/well of 3% non-fat, skim milk in PBST. Samples (serum or plasma) were diluted 1:100 in PBST; 100μL/well was added in duplicates and incubated at 37°C for 30 min. The excess antibody was washed off with PBST. Horseradish peroxidase (HRP) labeled-mouse anti-human IgG-Fc specific (GenScript No. A01854) diluted 1:10,000 in PBST was added (100μl/well) and incubated for 30 min at 37°C. After another washing step, the substrate solution (Sigma Aldrich No. P4809) was added (100μl/well) followed by 15 min incubation in dark. The reaction was stopped by the addition of 50μl/well 10% HCl and the absorbance was measured at 492nm (A_492_) using a Multiskan FC reader (Thermo Fisher Scientific). In every CovIgG-Assay determination two in-house controls, a high positive control (HPC) and negative control (NC) were included. HPC and NC were prepared by diluting an IgG anti-SARS-CoV-2 at a concentration of 30μg/ml and 0.070μg/ml, respectively in PBST containing 10% glycerol. The IgG anti-SARS-CoV-2 was purified from plasma of a convalescent patient using a 5/5 HiTrap rProtein-A column (GE Healthcare, USA). See detailed information about this procedure in **Supplementary method No.1**.

### Antibody class specificity

To confirm that our assay accurately detects antibody IgG class and excludes the potential for human IgM to cross-react with IgG, five (5) COVID-19 samples (1:100 diluted) were treated with 5mM DTT for 30 min at 37°C prior testing. After treatment, samples were added in duplicate (100μl/well) followed by the addition of the anti-human IgG-Fc-HRP (GenScript No. A01854) conjugate (diluted 1:10,000) or the addition of an anti-human IgM-HRP conjugate (Abcam No. ab97205) diluted 1:8,000 in PBST and the assay progressed as described above.

### Estimation of Antibody Titer

To estimate the IgG antibody titer, 40 COVID-19 samples were subjected to serial dilutions from 1:100 to 1:12,800. Each dilution was tested in duplicate in the CovIgG-Assay and each experiment was replicated twice. A standard curve was created in which the mean individual absorbance (A_492_) of each sample at 1:100 dilutions was correlated with its corresponding IgG antibody titer. Antibody titer was defined as the highest serum dilution that renders A_492_ values greater than the cut point estimated by the ROC analysis.

### Comparison with other serological assays approved for emergency use

We tested a set of 9 samples reported as IgM/IgG positives and 18 reported as IgG positives for SARS-CoV-2 antibodies by CoronaCheck (20/20 BioResponse, 20/20 Genesystems, Inc, Rockville, MA, USA). The information provided by the manufacturer claims that this assay use Roche’s technology (Roche Diagnostics GmbH, Sandhofer Strasse 116, D-68305 Mannheim, Germany). Same set of 18 samples reported as SARS-CoV-2 IgG positive were also tested by Abbott Architect SARS-CoV-2 IgG (Abbott Laboratories Diagnostics Division Abbott Park, IL 60064 USA). For comparison, both set of samples (n=27) were tested with our CovIgG-Assay. Moreover, another set of 18 samples from convalescent COVID-19 subjects, which had been confirmed by PCR were tested by CovIgG-Assay and Elecsys Anti-SARS-CoV-2 method (Cobas).

### Data analysis

Each CovIgG-Assay determination was performed in duplicate and the results expressed as the mean absorbance at 492 nm (A_492_) for each determination. The optimal cut point for the assay was established within a 95% confidence interval (CI) by receiver operating characteristic (ROC) curve analysis using the EpiTools epidemiological calculator (http://epitools.ausvet.com.au). Arbitrary guidelines were followed for analyzing the area under curve (AUC) as follows: non-informative, AUC=0.5; low accurate, 0.5 < AUC < 0.7; moderately accurate, 0.7 < AUC < 1; perfect, AUC = 1 (9). Intra-plate repeatability was evaluated for CovIgG-Assay by measuring the coefficient variation (CV) of 60 repeats of a High Positive Control (HPC) and a Negative Control (NC). For reproducibility evaluation, we completed three independent runs in different days for the CovIgG Assay using HPC, NC (30 replicates), four (4) negative and four (4) COVID-19 positive sera (6 replicates). Correlation between the A_492_ at 1:100 dilutions and the antibody titer as well as between the results of CovIgG Assay and the RT-PCR test results were evaluated using the Pearson correlation coefficient (with the 95% CI). To evaluate the agreement between the CovIgG-Assay and the RT-PCR, CovIgG-Assay and CoronaCheck, Abbott Architect SARS-CoV-2 IgG or Elecsys, inter-rater agreement (kappa) was calculated according to the method described by Thrusfield (10). The Kappa values (κ) were considered as follows: poor agreement, κ<0.02); fair agreement, κ=0.21 to 0.4; moderate agreement, κ=0.41 to 0.6; substantial agreement, κ=0.61 to 0.8; very good agreement, κ=0.81 to 1.0.

## Results

### Distribution of absorbance values of sera and ROC analysis

We used the RT-PCR for COVID-19 positive samples, as recommended standard reference method, to build ROC curves on the basis of the absorbance values (A_492_) obtained with specimens from two reference populations: subjects infected with SARS-CoV-2 that were all RT-PCR positive (assumed infected population) and healthy subjects or subjects that had been diagnosed with other respiratory or viral infections prior to the COVID-19 pandemic (uninfected population). The A_492_ values of uninfected population ranged between 0.011 and 0.312 with a mean ± SD A_492_ value of 0.075 ± 0.052 whereas samples from assumed infected population showed A_492_ values that ranged between 0.045 (one sample) and 3.21 with a mean A_492_ value of 1.99 ± 0.727. The mean value of the infected population was significantly different from the mean value of the uninfected population (*p*<0.0001). The distribution of A_492_ values of these two reference populations was very different. Approximately the 75% of infected population had A_492_ values between 0.828 and 2.5 (median 2.01), whereas that the 95% of uninfected population had A_492_ values between 0.011 and 0.176 (median 0.065) (**Figure-1**). Receiving operating characteristic analysis was used to determine the best cut-points for the CovIgG-Assay. The ROC optimized cut-point was 0.312. The selection of this cut-point derived from three different conditions: (a) maximum specificity at which the sensitivity was still 100%, (b) maximum sensitivity at which the specificity of the assay was also maximized, and (c) maximum value for Youden’s J index (S + Sp-1) and test efficiency (**Table-1**).

**Figure 1:**
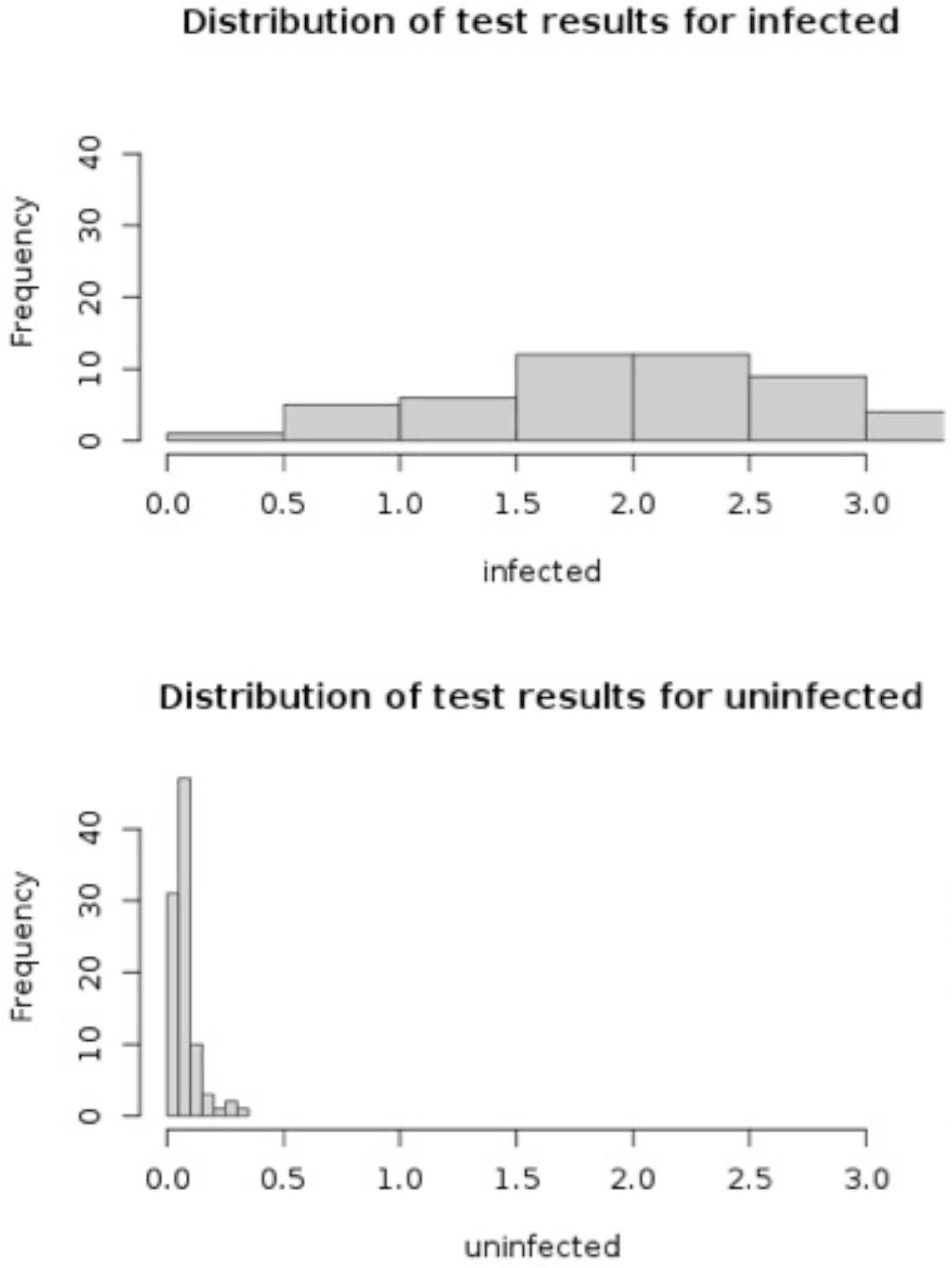
Distribution of absorbance values obtained with CovIgG-Assay. The CovIgG-Assay was optimized using as antigen recombinant Spike-S1-RBD from SARS-CoV-2. Absorbance values were distributed in form of frequency histograms to clearly visualize the separation between true positives (SARS-CoV-2 infected) (upper figure) and true negative population (non-SARS-CoV-2 infection), which include healthy subjects and subjects carrying other respiratory and viral infections collected prior pandemic (lower figure).

**Table-1.**
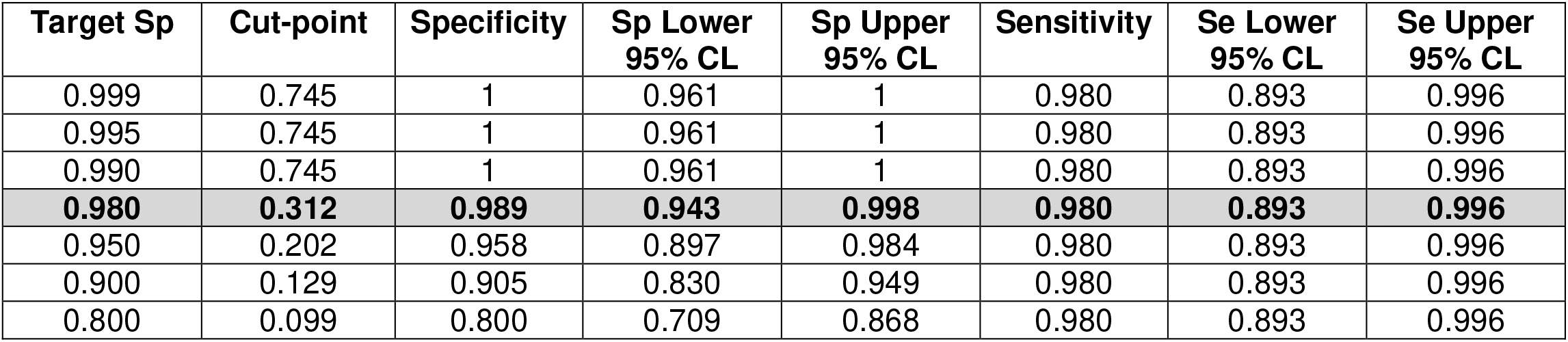
Specificity (Sp) and sensitivity (Se) of the CovIgG-Assay based on cut-points from the comparison between infected and uninfected populations.

The area under curve values (AUC) (accuracy) for the ROC curve was 0.985 (**Figure-2**). Based on the established cut-point only one seronegative was detected in the infected group whereas no seropositive was detected in the uninfected group. A sample from the uninfected group, collected between 1995 and June 2019, had A_492_ values equal to the cut-point and was considered negative. Significant differences (*p*<0.0001) were obtained between the mean OD values of COVID-19 infection sera (1.99 ± 0.727) compared to those from uninfected subjects (0.075 ± 0.052).

**Figure 2:**
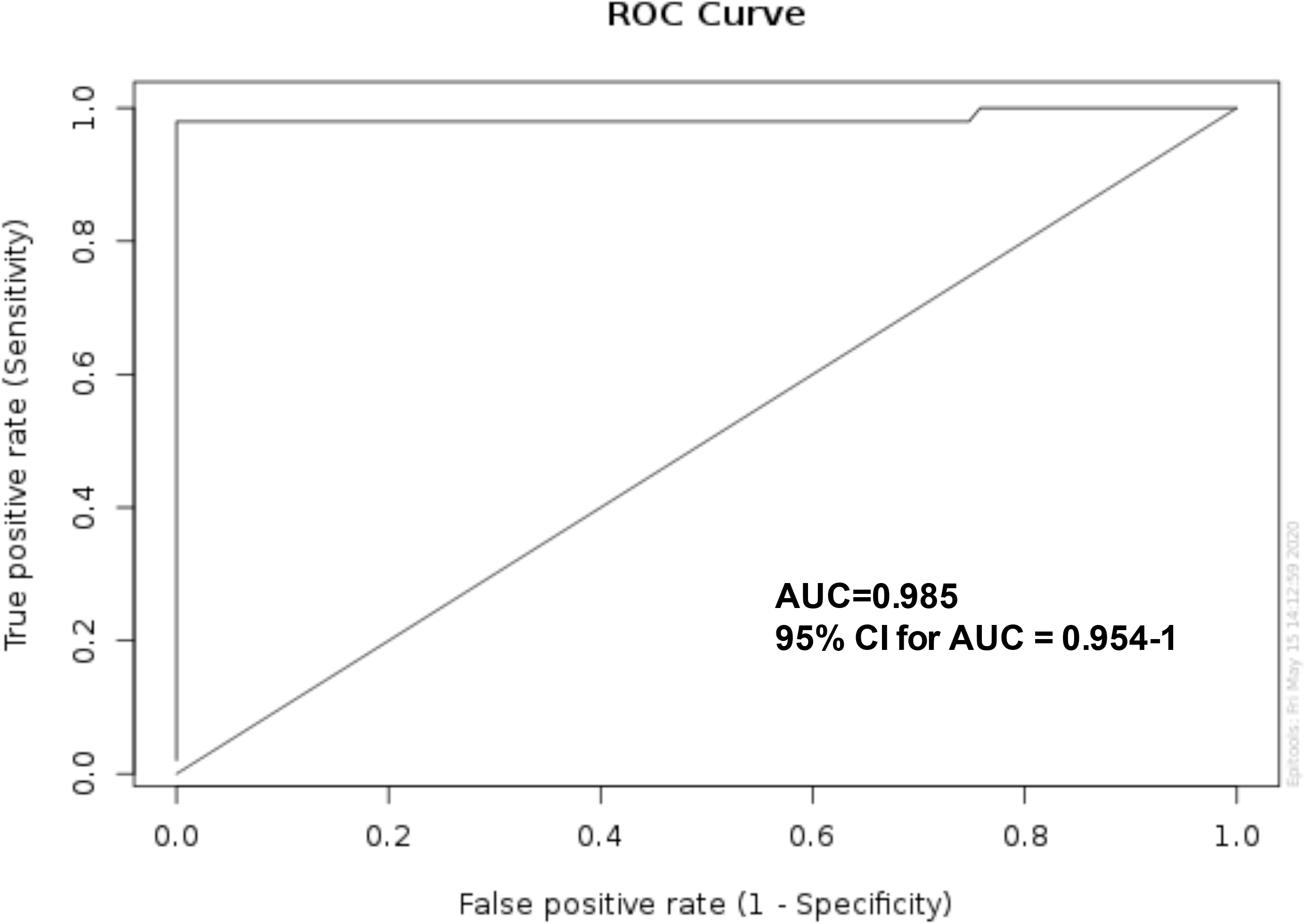
Receiver-operator characteristic (ROC) curve. The ROC curve was built for 132 sera from healthy subjects or subjects carrying other respiratory or other viral infections and 49 COVID-19 confirmed subjects. The area under the ROC curve (accuracy) was 0.985 and and the 95% Confident interval (CI) for AUC= 0.954-1.

To verify the cross reactivity of the assay we tested 67 samples know to be positive to common respiratory and non-respiratory infections (RSV, Flu A and B, Zika, and dengue) or allergies which are very common in the local population and that had been collected prior pandemic. As it is showed in figure 3, all those samples were negative showing no-cross reactivity in CovIgG-Assay. Thus, under these optimized conditions, CovIgG-Assay reached 98.9% specificity and 98.0% sensitivity with estimated predictive positive value (PPV) and predictive negative value (PNV) for CovIgG-Assay of 100% and 99.2%, respectively (**Table-2**). Importantly, all the 12 samples from individuals with Mycoplasma and Chikungunya that were collected during pandemic also resulted negative in the CovIgG-Assay (Figure-3), which confirmed the absence of cross-reaction in the CovIgG-Assay. There was substantial agreement (97.95%, κ=0.657) between CovIgG-Assay and RT-PCR. Detailed optical densities (ODs) values of the positive and negative samples, including the cross-reactivity panel are provided (**Supplementary tables 1 and 2 respectively**). We also assessed the reproducibility of the CovIgG-Assay by calculating the CV of data from 3 different assays and 30 repeats of controls and 6 repeats of selected negative and positive samples. The intra-assay and inter-assay reproducibility values were both lower than 10% (**Supplementary Table 3**).

**Figure 3:**
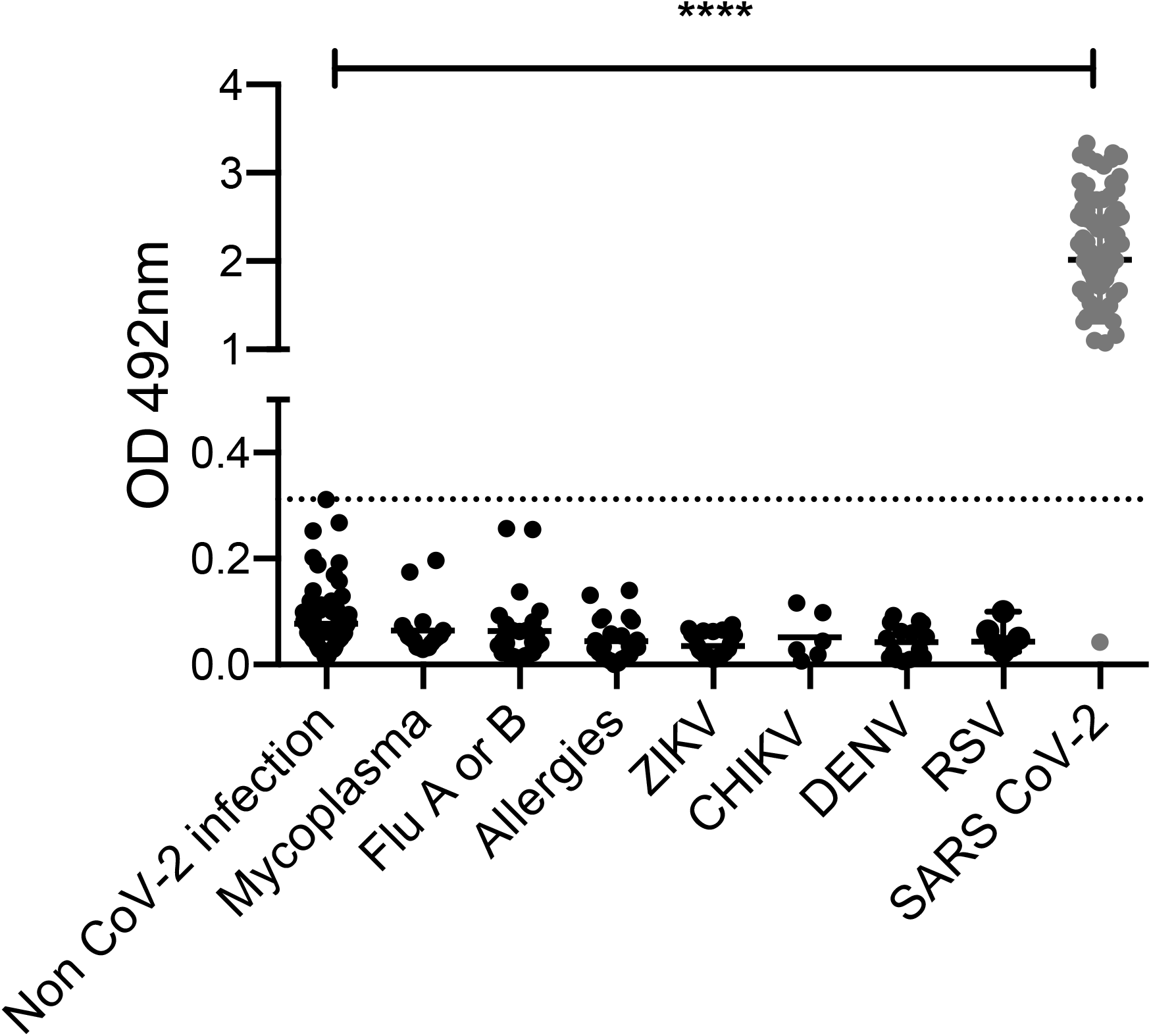
Validation of use of Spike S1-RBD ELISA for detection of SARS-CoV-2 IgG antibodies. Black dots indicate samples from negative cohorts (non-CoV-2 infection) collected prior 2019 with no previous history of selected viral infections or respiratory allergies (n= 78) and samples that tested positive for Mycoplasma IgM, (n = 9), Influenza A or B (n=13), respiratory allergies (n=13), Zika virus (ZIKV, n=14), Chikungunya virus (CHIKV, n=3), Dengue virus (DENV, n=8) or RSV (n=6). Grey dots indicate samples from patients with confirmed SARS CoV-2 infection (n=49). Dotted horizontal line indicate CovIgG-Assay cut-point value (OD_492_= 0.312). S1-RBD: Spike subunit-1-Receptor biding domaine. Each dot indicates mean OD of each sample tested in duplicate.

**Table-2.**
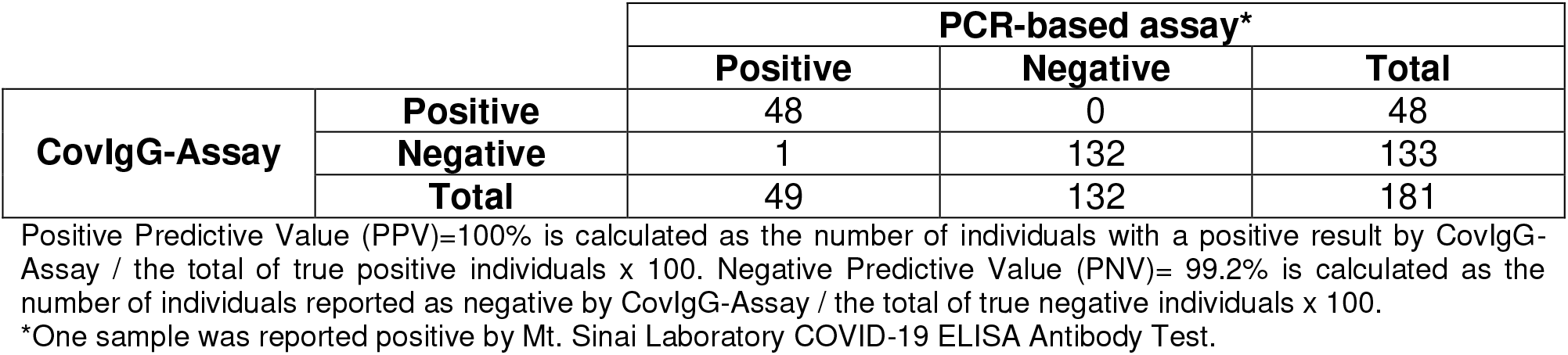
Agreement between the results of CovIgG-Assay and the PCR-based assay used as reference method for COVID-19 diagnosis.

### Class antibody specificity of CovIgG-Assay

To confirm that the positivity showed by CovIgG-Assay with the COVID-19 samples was mostly due to the presence of IgG antibody class and not due to potential cross-reactions with IgM antibody, five samples treated with DTT were tested in parallel on the CovIgG-Assay using as secondary antibody anti-human IgG- and anti-human IgM-HRP conjugates and the results obtained were compared with those obtained for the same samples previous to the DTT treatment. As expected, the A_492_ values of DTT-treated samples tested with the anti-IgM-HRP conjugate significantly dropped to values similar to the background. In contrast, the A_492_ values for the same DTT-treated samples tested with the anti-IgG-HRP conjugate were similar to those obtained with the untreated samples, confirming that positive results were from IgG antibodies. (**Table-3**).

**Table-3.**
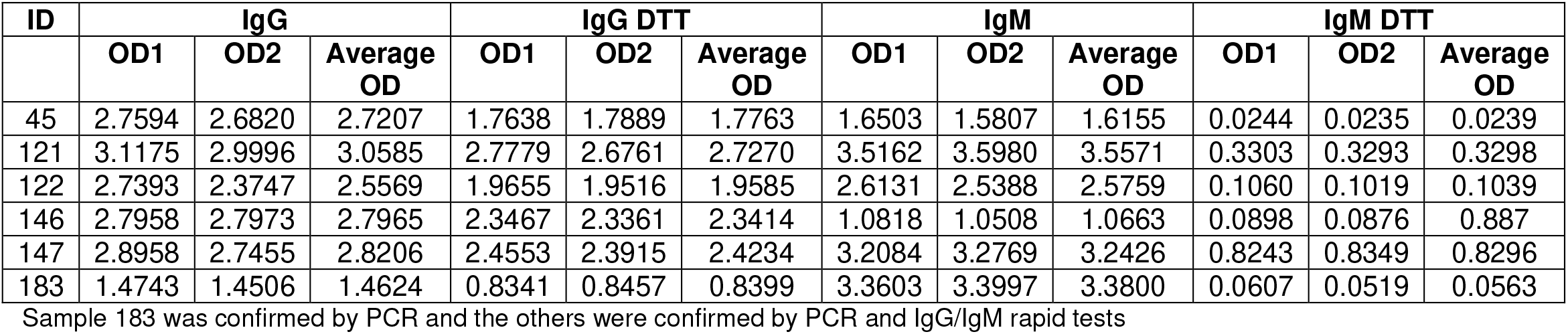
Samples before and after treatment with DTT (Dithiothreitol) showing IgG class specificity for CovIgG Assay. Each result represents OD at 492nm absorbance.

| ID | IgG |  |  | IgG DTT |  |  | IgM |  |  | IgM DTT |  |  |
| --- | --- | --- | --- | --- | --- | --- | --- | --- | --- | --- | --- | --- |
|  | OD1 | OD2 | Average OD | OD1 | OD2 | Average OD | OD1 | OD2 | Average OD | OD1 | OD2 | Average OD |
| 45 | 2.7594 | 2.6820 | 2.7207 | 1.7638 | 1.7889 | 1.7763 | 1.6503 | 1.5807 | 1.6155 | 0.0244 | 0.0235 | 0.0239 |
| 121 | 3.1175 | 2.9996 | 3.0585 | 2.7779 | 2.6761 | 2.7270 | 3.5162 | 3.5980 | 3.5571 | 0.3303 | 0.3293 | 0.3298 |
| 122 | 2.7393 | 2.3747 | 2.5569 | 1.9655 | 1.9516 | 1.9585 | 2.6131 | 2.5388 | 2.5759 | 0.1060 | 0.1019 | 0.1039 |
| 146 | 2.7958 | 2.7973 | 2.7965 | 2.3467 | 2.3361 | 2.3414 | 1.0818 | 1.0508 | 1.0663 | 0.0898 | 0.0876 | 0.887 |
| 147 | 2.8958 | 2.7455 | 2.8206 | 2.4553 | 2.3915 | 2.4234 | 3.2084 | 3.2769 | 3.2426 | 0.8243 | 0.8349 | 0.8296 |
| 183 | 1.4743 | 1.4506 | 1.4624 | 0.8341 | 0.8457 | 0.8399 | 3.3603 | 3.3997 | 3.3800 | 0.0607 | 0.0519 | 0.0563 |

Sample 183 was confirmed by PCR and the others were confirmed by PCR and IgG/IgM rapid tests

### Correlation between the A_492_ values and the IgG antibody titer

To determine whether the magnitude of the A_492_ values correlate with the antibody titer we selected 40 samples from infected individuals with A_492_ values among 0.321 to 3.12, which were the lowest and the greatest A_492_ obtained from the sample population studied, respectively. All 40 sera were diluted from 1:100 to 1:12,800 and each dilution was tested in duplicate in the CovIgG-Assay. The number of individuals with different antibody titers (defined as the maximal dilution that renders a positive result) is shown in Table-4. We found a lineal correlation (r^2^=0.9946) between the antibody titer (maximal dilution that render A_492_ > 0.312) and the individual A_492_ value at the working dilution (1:100). Thus, results reported by CovIgG-Assay could be quantitatively reported by estimating the titer, using the lineal equation (Y= 1.268*X −2.036) derived from the lineal correlation between antibody titer and the magnitude of absorbance values (**Figure-4**). Based on this analysis antibody estimated titers are reported in the range among 1:100 to 1:12,800. Samples with A_492_ in the range of 0.312 to 0.49 would have antibody titer lower than 1:100. Such a samples would be considered as weakly positive with undetermined antibody titer. It would be highly recommendable that another sample from such subjects collected 2-3 weeks thereafter can be tested. Samples with A_492_ >3.12 are reported with estimated antibody titer >1:12,800 (**Supplementary table 4**).

**Figure 4:**
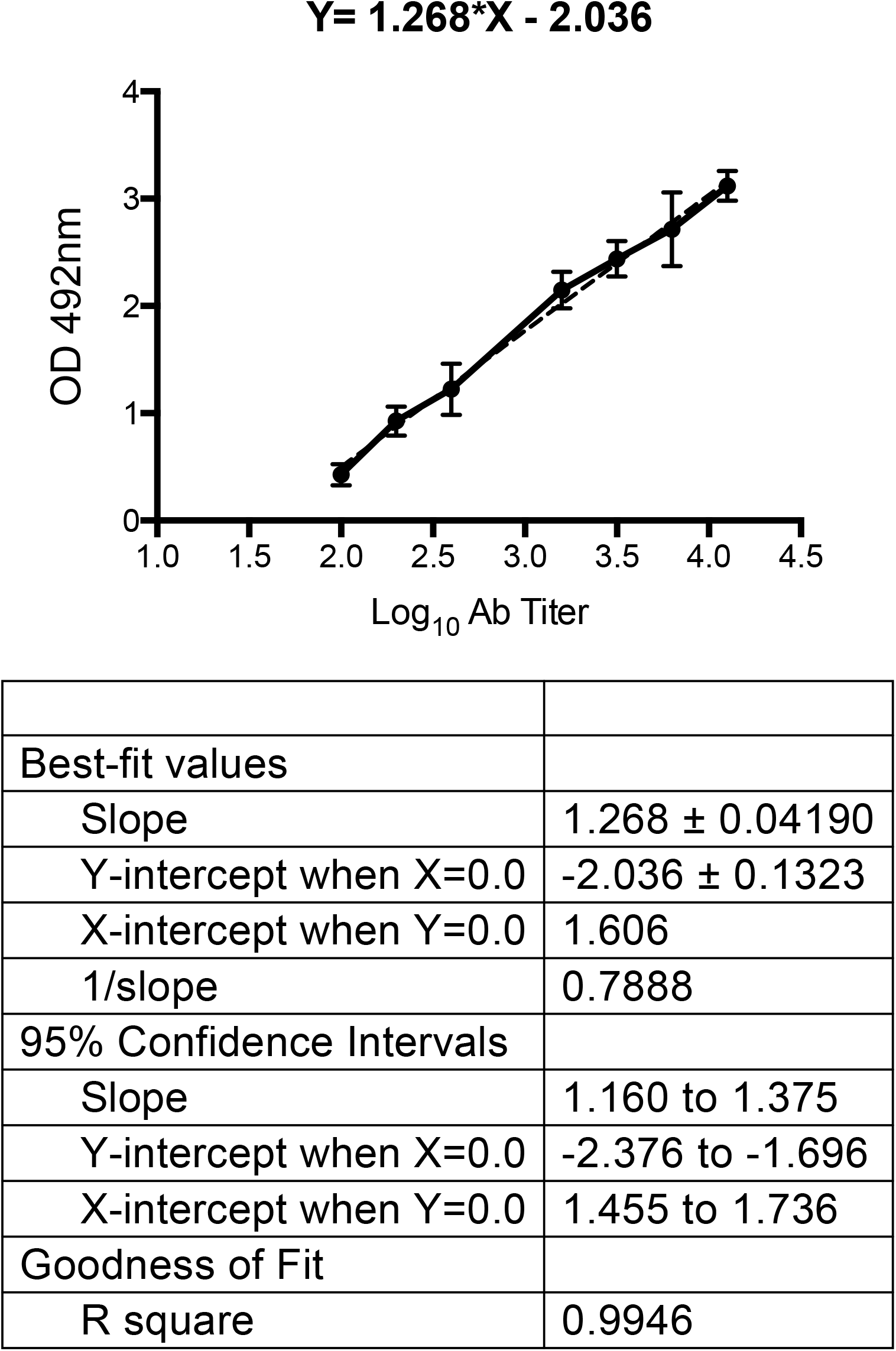
Correlation between absorbance at 492nm (A_492_) and antibody titter. A total of 40 sera from confirmed COVID-19 subjects that resulted positive by CovIgG-Assay were titrated at dilutions among 1:100 to 1:12,800. A lineal regression analysis was then done in which the mean A_492_ of sera with similar antibody titer were plotted with their corresponding A_492_ values. We found a lineal correlation (r2=0.9946) between the antibody titer (maximal dilution that render A_492_ ≥ 0.312) and the individual A_492_ value at the working dilution (1:100). From this analysis the following lineal equation (Y= 1.268*X −2.036) was obtained, which was further used to estimate the antibody titer of all sera reported seropositive by CovIgG-Assay.

**Table-4.** Experimental antibody titers of SARS-CoV-2 infected subjects in relation with their absorbance value at 492nm (1:100 dilution).

| **Antibody titer\*** | **No. Individuals** | **Absorbance at 492nm (A<sub>492</sub>) range** | **Mean A<sub>492</sub> ± SD** |
| --- | --- | --- | --- |
| 100 | 6 | 0.321-0.538 | 0.427 ± 0.0983 |
| 200 | 10 | 0.687-1.111 | 0.927 ± 0.135 |
| 400 | 8 | 0.802-1.518 | 1.224 ± 0.238 |
| 800 | -- | -- | -- |
| 1600 | 6 | 1.980-2.385 | 2.149 ± 0.169 |
| 3200 | 6 | 2.175-2.577 | 2.439 ± 0.165 |
| 6400 | 2 | 2.372-3.059 | 2.715 ± 0.343 |
| 12800 | 2 | 2.851-3.128 | 2.985 ± 0.138 |

\*Antibody titer is defined as the maximal serum dilution that renders A<sub>492</sub> greater than the optimized cut-point (A<sub>492</sub> ≥ 0.312) determined by ROC analysis.

### Agreement between CovIgG-Assay and other serological tests

To evaluate the performance of CovIgG-Assay with other tests in the market, we analyzed a group of samples that have been previously reported as positive for IgG/IgM (n=9) or only positive for IgG (n=18) by CoronaCheck rapid test. CovIgG-Assay had 100% agreement with the CoronaCheck results for the IgG/IgM positive samples. These samples were all reported as positive by CovIgG-Assay with antibody titers that ranged between 1:100 and 1:3,251. However, the agreement was fair (38.88%) for samples only reported positive for IgG by CoronaCheck since 7 from 18 samples were reported as positive by CovIgG-Assay (**Table-5**). Interestingly, all 18 these presumptive IgG positive samples were found negative by the Abbot Architec SARS-CoV-2 IgG method, which reveal a better agreement (61.0%) between our CovIgG-Assay method and Abbot Architec SARS-CoV-2 IgG (**Table-5**) and might suggest that most of these 18 samples could be false positives.

**Table 5.** Comparison of three serological tests against CoVIgG-Assay for detection of anti SARS CoV-2 antibodies.

| Sample ID | CoronaCheck<sup>1</sup><br>(Rapid COVID-19 IgG/IgM) | CoronaCheck<sup>1</sup><br>(Rapid COVID-19 IgG) | CoVlgG-Assay<sup>2</sup><br>Estimated titer | Abbott Architect SARS-CoV-2 IgG<sup>3</sup><br>(Index value) | Elecsys<sup>4</sup><br>Anti SARS CoV-2<br>Cobas® |
| --- | --- | --- | --- | --- | --- |
| 32 | + |  | >12800 |  |  |
| 33 | + |  | >12800 |  |  |
| 34 | + |  | 1:649 |  |  |
| 35 | + |  | 1:11040 |  |  |
| 36 | + |  | 1:1963 |  |  |
| 37 | + |  | >12800 |  |  |
| 38 | + |  | 1:8241 |  |  |
| 39 | + |  | >12800 |  |  |
| 40 | + |  | >12800 |  |  |
| 46 |  | + | - | .05 |  |
| 47 |  | + | † | .03 |  |
| 48 |  | + | - | .03 |  |
| 49 |  | + | 1:254 | .05 |  |
| 50 |  | + | - | .03 |  |
| 51 |  | + | - | .05 |  |
| 52 |  | + | - | .02 |  |
| 53 |  | + | - | .04 |  |
| 54 |  | + | 1:254 | .05 |  |
| 55 |  | + | - | .04 |  |
| 56 |  | + | - | .07 |  |
| 57 |  | + | - | .37 |  |
| 58 |  | + | † | .08 |  |
| 59 |  | + | † | .02 |  |
| 60 |  | + | - | .05 |  |
| 61 |  | + | - | .01 |  |
| 62 |  | + | 1:219 | .69 |  |
| 63 |  | + | 1:220 | .05 |  |
| 105 |  |  | 1:3200<sup>‡</sup> |  | + |
| 106 |  |  | 1:200<sup>‡</sup> |  | + |
| 132 |  |  | 1:3200<sup>‡</sup> |  | + |
| 133 |  |  | 1:400<sup>‡</sup> |  | + |
| 134 |  |  | 1:200<sup>‡</sup> |  | - |
| 154 |  |  | 1:200<sup>‡</sup> |  | + |
| 148 |  |  | 1:1600<sup>‡</sup> |  | + |
| 149 |  |  | 1:800<sup>‡</sup> |  | + |
| 150 |  |  | 1:200<sup>‡</sup> |  | + |
| 151 |  |  | 1:1600<sup>‡</sup> |  | + |
| 152 |  |  | 1:400<sup>‡</sup> |  | + |
| 153 |  |  | 1:6400<sup>‡</sup> |  | + |
| 194 |  |  | >1:12800 |  | + |
| 195 |  |  | >1:12800 |  | + |
| 197 |  |  | 1:1064 |  | + |
| 220 |  |  | >1:12800 |  | + |
| 221 |  |  | >1:12800 |  | + |
| 222 |  |  | 1:3236 |  | + |

<sup>‡</sup>Titer determined by endpoint dilution from the average of duplicates OD492.
<sup>1</sup>Lateral flow rapid test intended to qualitatively detect antibodies to SARS-CoV-2. Presence of IgG/IgM antibodies are indicated by a visible line in the specific region on the device (On the manufacturer's website mention holding FDA EUA). Positive results may be due to cross reactivity with other Coronavirus strains (non CoV-2). Information regarding antigen used is not available.
<sup>2</sup>Quantitative ELISA for the detection SARS CoV-2 IgG antibodies using Spike S1-RBD (GenScript) as capture antigen. Samples with OD492<0.312 are considered negative. †IgG titration of positive samples by CovIgG-Assay range from 1:100 to 1:12,800.
<sup>3</sup>Chemiluminescent microparticle immunoassay for the qualitative detection of IgG antibodies against SARS CoV-2 (FDA EUA).
Assay is designed to detect IgG antibodies to the nucleocapsid protein of SARS CoV-2. Samples with an index value <1.4 are considered negative. <sup>4</sup>Electrochemiluminescence immunoassay intended for qualitative detection of antibodies to SARS-CoV-2 in human serum and plasma using a recombinant protein representing the nucleocapsid antigen of SARS-CoV-2 (FDA EUA).

In another experiment, samples from subjects confirmed by PCR (n=18) were analyzed by our CovIgG-Assay and Elecsys Anti-SARS-CoV-2 method. CovIgG-Assay reported as positive all these 18 samples with antibody estimated titers ranging among 1:200 and >1:12,800 whereas Elecsys Anti-SARS-CoV-2 method reported 17 positive. Thus, very good agreement (94.4%) between Elecsys and our CovIgG-Assay was observed (**Table-5**).

## Discussion

Since the circulation of SARS-CoV-2 was spillover outside of China, the global efforts to develop serological assays have been unprecedently huge. The precise diagnostic of COVID-19 poses multiples challenges. The proposed method reference is the molecular diagnostic, which determines the presence of an active infection. However the timing of viral replication and the development of immune response is quite variable (reviewed in (11) and the presence of IgM and or IgG at the time of the molecular diagnostic is merely speculative. While the molecular testing results are a guide, they should not be considered gold standards, as it has been the practice so far. Other factors as clinical presentation and epidemiological aspects need to be considered at the time of selecting the appropriate samples to validate any assay. Here we selected 48 samples reported positive by authorized molecular methods and 1 sample reported as positive by an authorized serological assay which is not considered a rapid test (7). As negative samples we used a set of 132 sera that were collected in the period of 1995 to June 2019, before COVID-19 period. The CovIgG-Assay data were subjected to ROC curve analysis. During the last two decades this type of analysis has become a popular method for evaluating the accuracy of medical diagnostic systems and has been used not only to evaluate the ability of a test to discriminate between infected and healthy subjects (12) but also to compare the diagnostic performance of a number of tests (13). The ROC curve is obtained by plotting the true-positive rate (sensitivity) as a function of the false-positive rate (100-specificity) that is associated with each cut-point. The AUC is then used as a measure of the accuracy of the test. If the assay can distinguish between infected and normal populations the AUC will be equal to 1 and ROC curve will reach the upper left corner. As our results demonstrate, the AUC value obtained from the ROC curve analysis conducted on the CovIgG-Assay data was very high, indicating the high accuracy of this test. The only sample that was not detected by CovIgG-Assay could be a false positive in the RT-PCR. Currently, there are few reports addressing those discordant results. Several molecular assays have been developed with high specificity and low limit of detections (14–17) and are considered the reference method for SARS-CoV-2 diagnostic (18). However everyday there are more reports addressing problems with the RT-PCR accuracy (14, 19–21). Otherwise that sample may be collected within a window where the immune response was not developed yet. Nevertheless, our results reinforce the complexity of the diagnostic of COVID-19 and the need for prospective studies with more samples and better-characterized cohorts to expand our understanding of the dynamics of the immune response to this novel coronavirus.

We also considered fundamental to develop a quantitative test, in addition to suggesting a qualitative result. This would provide a guide about quantity or the dimension of the immune response mounted by an individual. Up to today, few works on COVID-19 addressed the relation between the titer of the IgG and the neutralizing capability of that sample. But all of them coincide that there is a direct correlation (1, 21–23). While the scientific community develops safe (BSL-2) and reproducible neutralization assays to determine nAbs against SARS-CoV-2, quantitative assays like CovIgG-Assay are useful tools for a reliable serological characterization.

The notable disagreement observed between CoronaCheck and Abbot Architec SARS-CoV-2 IgG method for the same set of samples (100% and 0% positive samples, respectively) resulted surprising and at the same time worrisome since both methods have FDA EUA. Since these samples were kindly donated by Clinical Laboratories de-identified we unknown whether the presence of virus was confirmed in any of these subjects. However, because our assay had significant agreement with the Abbot method (61%) and very good agreement (94%) with Elecsys method using a set of samples from PCR-confirmed SARS-CoV-2 infection, we could suggest that most IgG positive samples reported by CoronaCheck might be false positive. Furthermore we have recent results showing that the sample reported as negative by Elecsys and positive by CovIgG-Assay had neutralizing antibodies (data not shown-manuscript in preparation). Together those results reinforce the notion that our method is capable to offer accurate and efficient diagnostic testing for detection of antiviral antibodies in infected individuals.

The finding of contradictory results in the performance of different immunoenzymatic diagnostic methods strengthen the need for better assays and for better validations in the context of clinical presentations (24). The best characteristics of CovIgG-Assay are its PPV of 100% along a PNV of 99.2%, which render highly trustable positive results. A subject confirmed as positive by this assay, with a high degree of confidence can be reincorporating to a normal life in support of the rest of the community. On the other hand the high NPV of the assay increase its value to know which individuals may be still at risk to be infected. Altogether, the quantification of the IgG, after a subject being reported as positive, provides a second step of certainty of possible protection against SARS-CoV-2 infection. However such correlation studies using a larger set of samples are under way.

## Authors Contribution

AME and CAS conceived and supervised the studies. PP and AME performed the experiments. CAS and AME drafted the manuscript. CAS, AME and PP reviewed the final version of the manuscript. CAS and AME obtained the funds. CAS prepared and CAS and AME submitted the Emergency Use Authorization request to the US Federal Drug Administration.

## Acknowledgement

Authors want to thank Ilia Toledo, MT, Francheska Rivera, MT and Drs. Consuelo Climent, Gerardo Latoni and Ivelisse Martin for their contribution and diligent access to the samples from subject presumptive exposed to SARS-CoV-2 and from plasma donors. Also, thanks to Dr. Jorge L. Muñoz-Jordan for providing the panel to improve the cross-reactivity testing. Particular acknowledgement deserves all administrative and supportive staff at the Medical Sciences Campus, University of Puerto Rico, Laboratorio Clinico Toledo, Laboratorio Clinico Martin, Banco de Sangre Centro Médico and Banco de Sangre Servicios Mutuos for their availability and commitment during the curfew imposed by the quarantine period. The Puerto Rico Science, Technology and Research Trust supported research reported in this work under agreement number 2020-00272 to AME and CAS. The excellent support of Ms. Andreica Maldonado and Grace Rendon from PRSTRT was instrumental in support of the work described here.

## SUPPLEMENTARY DATA

### Supplementary Method-1

#### IgG Purification for positive and negative controls elaboration

For the assay quality control; positive and negative controls were included for each assay. These controls were prepared in-house from a convalescent subject with COVID-19. **(A)** 20mL of the plasma sample was mixed 1:1 with phosphate buffer saline (PBS) and loaded onto a 5/5 HiTrap rProtein-A column (GE Healthcare, USA) at a flow rate: of 1.5mL/min. The equilibration buffer used was PBS (pH 7.2), for elution we used 0.1M Glycine-HCl (pH 2.85) and the neutralization buffer used was 1M Tris pH 8.5. **(B)** A total of 17 fractions were collected and absorbance (OD) at 280nm for each fraction was measured. Fractions 9 to 14 were pooled in a total volume of 20ml and the pooled sample had A_280_=2.179. Pooled sample was desalted by PD-10 column and the desalted fraction had A_280_= 2.076). The final concentration for this fraction was 1.48mg/mL (2.076 /14) × 10) in 28mL, for a total of 41.44 mg. **(C)** A total of 2μg-purified IgG was suspended in loading SDS-buffer containing 5mM DTT and analyzed for purity by 4-20% sodium dodecyl sulfate polyacrylamide electrophoresis (SDS-PAGE) and stained with Coomassie-blue. The purified IgG fraction was used to prepare a high positive control (HPC) and a negative control (NC) to be used in the CovIgG-Assay. For the HPC preparation IgG was diluted 30μg/mL and for the NC IgG was diluted 380-fold to get a concentration equivalent to 0.078μgmL, both controls were prepared in PBST containing 10% glycerol.

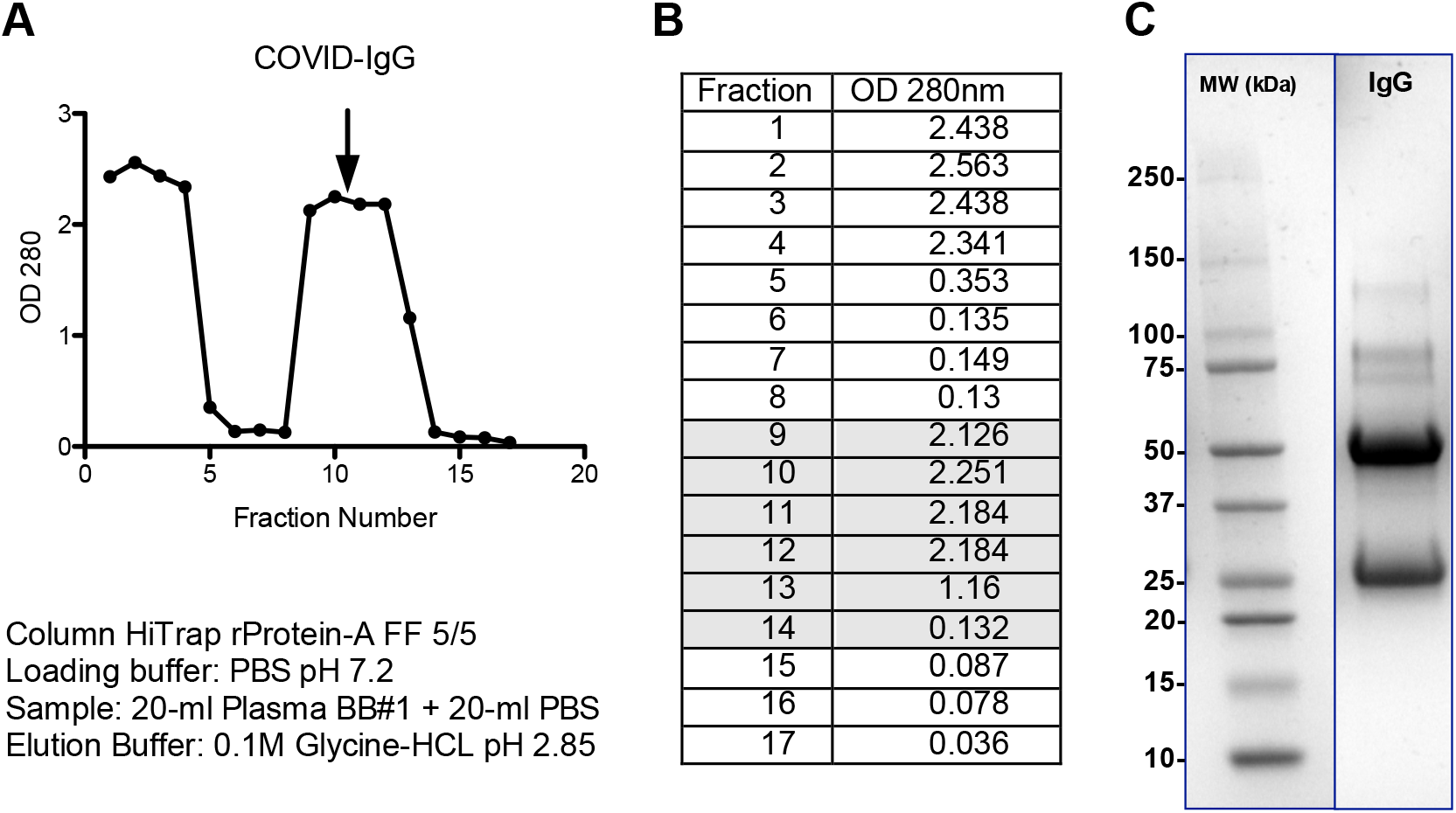

**Supplementary Table-1.** Positive samples used CovIgG-Assay validation. Each result represent the absorbance measurement at 492nm.

| Immunological status/Source | Numeric ID | Confirmatory Method | Sample Date | OD1 | OD2 | Average OD | Estimated Titer |
| --- | --- | --- | --- | --- | --- | --- | --- |
| PCR+/ IgG/IgM Positive Local Laboratory | 45 | LabCorp | 4/12/20 | 2.034 | 2.251 | 2.291 | 1:2691 |
| Blood Bank 1 PCR+ | 102 | Roche or LabCorp or CDC | 4/27/20 | 2.533 | 2.591 | 2.555 | 1:4487 |
|  | 103 | Roche or LabCorp or CDC | 4/27/20 | 2.584 | 2.463 | 2.539 | 1:4355 |
|  | 104 | Roche or LabCorp or CDC | 4/27/20 | 0.972 | 0.975 | 0.973 | 1:207 |
|  | 105 | Roche or LabCorp or CDC | 4/27/20 | 2.422 | 2.487 | 2.447 | 1:3639 |
|  | 106 | Roche or LabCorp or CDC | 4/27/20 | 0.797 | 0.876 | 0.826 | 1:156 |
| Local Major Hospital PCR+ | 120 | Cepheid | 4/27/20 | 2.022 | 2.005 | 2.014 | 1:1570 |
| Local Major Hospital PCR+ / IgG/IgM Positive | 121 | Cepheid | 4/24/20 | 2.390 | 2.355 | 2.372 | 1:3148 |
| Local Major Hospital PCR+ / IgG+/IgM- | 143 | Cepheid | 4/28/20 | 2.885 | 2.818 | 2.851 | 1:7980 |
| PCR+ / IgG/IgM Positive Local Laboratory | 122 | LabCorp | 4/16/20 | 1.991 | 1.970 | 1.981 | 1:1472 |
| Blood Bank 1 PCR+ | 132 | Roche or LabCorp or CDC | 4/30/20 | 2.153 | 2.198 | 2.176 | 1:2148 |
|  | 133 | Roche or LabCorp or CDC | 4/30/20 | 1.163 | 0.998 | 1.081 | 1:256 |
|  | 134 | Roche or LabCorp or CDC | 4/30/20 | 0.828 | 0.816 | 0.822 | 1:155 |
|  | 135 | Roche or LabCorp or CDC | 4/30/20 | 1.953 | 2.009 | 1.981 | 1:1472 |
|  | 136 | Roche or LabCorp or CDC | 4/30/20 | 0.048 | 0.043 | 0.045 | N/A |
|  | 137 | Roche or LabCorp or CDC | 4/30/20 | 0.973 | 1.100 | 1.037 | 1:235 |
| Blood Bank 2 PCR+ | 155 | Quest PCR | 5/11/20 | 1.429 | 1.314 | 1.371 | 1:450 |
|  | 156 | Quest PCR | 5/11/20 | 1.793 | 1.724 | 1.759 | 1:955 |
|  | 157 | Quest PCR | 5/11/20 | 2.464 | 2.501 | 2.483 | 1:3899 |
|  | 158 | Roche | 5/11/20 | 3.080 | 3.334 | 3.207 | >12800 |
|  | 159 | PR ASEM PCR | 5/11/20 | 2.906 | 3.206 | 3.056 | 1:11,885 |
|  | 160 | Quest PCR | 5/11/20 | 1.916 | 1.879 | 1.897 | 1:1,250 |
|  | 161 | Quest PCR | 5/11/20 | 2.746 | 3.170 | 2.958 | 1:9,817 |
|  | 162 | Roche | 5/11/20 | 2.264 | 1.981 | 2.122 | 1:1936 |
|  | 163 | Roche | 5/11/20 | 1.415 | 1.460 | 1.437 | 1:512 |
|  | 164 | Roche | 5/11/20 | 2.694 | 2.513 | 2.604 | 1:4932 |
|  | 165 | Roche | 5/11/20 | 2.199 | 2.194 | 2.196 | 1:2239 |
|  | 166 | Roche | 5/11/20 | 1.715 | 1.851 | 1.783 | 1:1002 |
|  | 167 | Roche | 5/11/20 | 1.796 | 1.800 | 1.798 | 1:1032 |
|  | 170 | VA Orlando, FL PCR | 5/11/20 | 3.128 | 3.152 | 3.140 | >12800 |
|  | 171 | Roche | 5/11/20 | 2.468 | 2.487 | 2.477 | 1:3864 |
|  | 173 | PR Dept. of Health PCR | 5/11/20 | 3.185 | 3.223 | 3.204 | >12800 |
|  | 176 | VA San Juan PCR | 5/11/20 | 2.484 | 2.541 | 2.512 | 1:4130 |
|  | 177 | PR Dept. of Health PCR | 5/11/20 | 2.103 | 2.296 | 2.200 | 1:2254 |
|  | 178 | Quest PCR | 5/11/20 | 1.621 | 1.521 | 1.571 | 1:664 |
|  | 181 | Roche | 5/11/20 | 1.748 | 1.682 | 1.715 | 1:879 |
|  | 182 | Quest PCR | 5/11/20 | 2.857 | 2.955 | 2.906 | 1:8872 |
|  | 183 | COVID-19 Ab Assay - Titer<br>2880 Mount Sinai | 5/11/20 | 1.758 | 1.663 | 1.711 | 1:871 |
|  | 184 | Quest PCR | 5/11/20 | 1.889 | 1.616 | 1.752 | 1:944 |
|  | 185 | Roche | 5/11/20 | 1.362 | 1.497 | 1.430 | 1:505 |
| **PCR+ Local**<br>**Laboratory** | 146 | LabCorp | 5/11/20 | 2.707 | 2.756 | 2.732 | 1:6324 |
|  | 147 | LabCorp | 5/11/20 | 2.573 | 2.700 | 2.636 | 1:5260 |
| **Blood Bank 1**<br>**PCR+** | 148 | Roche or LabCorp or CDC | 5/11/20 | 2.480 | 2.019 | 2.250 | 1:2483 |
|  | 149 | Roche or LabCorp or CDC | 5/11/20 | 2.052 | 2.131 | 2.092 | 1:1824 |
|  | 150 | Roche or LabCorp or CDC | 5/11/20 | 0.925 | 0.848 | 0.886 | 1:175 |
|  | 151 | Roche or LabCorp or CDC | 5/11/20 | 2.153 | 2.220 | 2.187 | 1:2193 |
|  | 152 | Roche or LabCorp or CDC | 5/11/20 | 1.316 | 1.073 | 1.195 | 1:319 |
|  | 153 | Roche or LabCorp or CDC | 5/11/20 | 2.365 | 2.155 | 2.260 | 1:2529 |
|  | 154 | Roche or LabCorp or CDC | 5/11/20 | 0.984 | 0.990 | 0.987 | 1:213 |

Estimated titer was calculated , using the lineal equation ( $Y = 1.185 \cdot X - 1.773$ ) derived from the lineal correlation between antibody titer and the magnitude of absorbance values.

**Supplementary Table 2.** Sera from subjects with non-SARS-CoV-2 infection used for CovIgG-Assay validation. Each result represent the absorbance measurement at 492nm.

| **Sample Source** | **Numeric ID** | **Immunological status** | **Sample Testing Date** | **OD1** | **OD2** | **Average OD** |
| --- | --- | --- | --- | --- | --- | --- |
| **Virology Serum Bank** | VB1 | **Healthy Subjects** | 5/23/95 | 0.028 | 0.027 | 0.027 |
|  | VB2 |  | 12/7/95 | 0.211 | 0.194 | 0.202 |
|  | VB3 |  | 2/10/97 | 0.082 | 0.087 | 0.085 |
|  | VB4 |  | 5/25/97 | 0.054 | 0.053 | 0.054 |
|  | VB5 |  | 2/17/06 | 0.041 | 0.054 | 0.047 |
|  | VB6 |  | 6/28/16 | 0.054 | 0.059 | 0.056 |
|  | VB7 |  | 4/10/17 | 0.092 | 0.106 | 0.099 |
|  | VB8 |  | 8/15/16 | 0.067 | 0.102 | 0.084 |
|  | VB9 |  | 8/15/16 | 0.310 | 0.225 | 0.268 |
|  | VB10 |  | 6/27/17 | 0.075 | 0.058 | 0.067 |
|  | VB11 |  | 1/10/17 | 0.023 | 0.000 | 0.011 |
|  | VB12 |  | 4/28/17 | 0.076 | 0.061 | 0.068 |
|  | VB13 |  | 6/26/18 | 0.330 | 0.294 | 0.312 |
|  | VB14 |  | 6/26/18 | 0.025 | 0.025 | 0.025 |
|  | VB15 |  | 6/28/18 | 0.036 | 0.030 | 0.033 |
|  | VB16 |  | 9/8/00 | 0.040 | 0.050 | 0.045 |
|  | VB17 |  | 5/8/19 | 0.026 | 0.033 | 0.029 |
|  | VB18 |  | 6/6/19 | 0.069 | 0.077 | 0.073 |
| **Immunology Bank** | IB95 | **Healthy Subjects** | 2012 | 0.087 | 0.094 | 0.090 |
|  | IB96 |  | 2012 | 0.042 | 0.050 | 0.046 |
|  | IB97 |  | 2012 | 0.081 | 0.076 | 0.078 |
|  | IB100 |  | 2012 | 0.032 | 0.031 | 0.032 |
|  | IB144 |  | 2012 | 0.100 | 0.100 | 0.100 |
|  | IB145 |  | 2012 | 0.048 | 0.036 | 0.042 |
|  | IB146 |  | 2012 | 0.072 | 0.067 | 0.069 |
|  | IB147 |  | 2012 | 0.035 | 0.038 | 0.036 |
|  | IB148 |  | 2012 | 0.028 | 0.031 | 0.029 |
|  | IB149 |  | 2012 | 0.043 | 0.049 | 0.046 |
|  | IB135 |  | 2012 | 0.044 | 0.044 | 0.044 |
|  | IB136 |  | 2012 | 0.112 | 0.095 | 0.103 |
|  | IB137 |  | 2012 | 0.024 | 0.027 | 0.025 |
|  | IB138 |  | 2012 | 0.023 | 0.034 | 0.029 |
|  | IB139 |  | 2012 | 0.028 | 0.027 | 0.027 |
|  | IB140 |  | 2012 | 0.107 | 0.106 | 0.106 |
|  | IB141 |  | 2012 | 0.033 | 0.025 | 0.029 |
|  | IB142 |  | 2012 | 0.049 | 0.040 | 0.044 |
|  | IB143 |  | 2012 | 0.041 | 0.040 | 0.040 |
|  | IM |  | 2012 | 0.032 | 0.029 | 0.030 |
|  | AO |  | 2012 | 0.096 | 0.090 | 0.093 |
|  | OF |  | 2012 | 0.075 | 0.069 | 0.072 |
|  | VB83 |  | 2012 | 0.064 | 0.060 | 0.062 |
|  | IB37 |  | 2012 | 0.069 | 0.078 | 0.074 |
|  | IB58 |  | 2012 | 0.055 | 0.059 | 0.057 |
|  | IB86 |  | 2012 | 0.016 | 0.015 | 0.015 |
|  | IB84 |  | 2012 | 0.014 | 0.016 | 0.015 |
| **Virology**<br>**Serum Bank** | RB | **ZIKV +** | 2016 | 0.084 | 0.082 | 0.083 |
|  | FM |  | 2016 | 0.019 | 0.024 | 0.021 |
|  | VB82 |  | 2016 | 0.083 | 0.088 | 0.085 |
|  | JN |  | 8/11/16 | 0.076 | 0.054 | 0.065 |
|  | JR |  | 8/15/16 | 0.157 | 0.157 | 0.157 |
|  | EXP |  | 4/1/17 | 0.040 | 0.015 | 0.028 |
| **Immunology**<br>**Bank** | IB133 | **Healthy Subjects** | 2012 | 0.029 | 0.004 | 0.017 |
|  | IB130 |  | 2012 | 0.073 | 0.061 | 0.067 |
|  | IB131 |  | 2012 | 0.024 | 0.025 | 0.024 |
|  | IB132 |  | 2012 | 0.253 | 0.251 | 0.252 |
| **Immunology**<br>**Bank** | IB1 | **Resp. Allergies** | 2012 | 0.081 | 0.084 | 0.083 |
|  | IB2 |  | 2012 | 0.044 | 0.048 | 0.046 |
|  | IB3 |  | 2012 | 0.048 | 0.049 | 0.049 |
|  | IB4 |  | 2012 | 0.055 | 0.123 | 0.089 |
|  | IB5 |  | 2012 | 0.127 | 0.061 | 0.094 |
|  | IB6 |  | 2012 | 0.050 | 0.050 | 0.050 |
|  | IB7 |  | 2012 | 0.103 | 0.107 | 0.105 |
|  | IB8 |  | 2012 | 0.081 | 0.080 | 0.081 |
|  | IB9 |  | 2012 | 0.090 | 0.095 | 0.093 |
|  | IB10 |  | 2012 | 0.003 | 0.045 | 0.024 |
|  | IB11 |  | 2012 | 0.255 | 0.128 | 0.192 |
|  | IB12 |  | 2012 | 0.065 | 0.064 | 0.065 |
|  | IB13 |  | 2012 | 0.058 | 0.317 | 0.188 |
| **Immunology**<br>**Bank** | BVF |  | 2012 | 0.052 | 0.059 | 0.056 |
|  | VA |  | 2012 | 0.058 | 0.064 | 0.061 |
|  | JJF |  | 2012 | 0.070 | 0.075 | 0.073 |
|  | IB53 |  | 2012 | 0.067 | 0.062 | 0.065 |
|  | IB54 |  | 2012 | 0.058 | 0.059 | 0.059 |
|  | IB55 |  | 2012 | 0.049 | 0.050 | 0.050 |
|  | IB56 | **Healthy Subjects** | 2012 | 0.180 | 0.158 | 0.169 |
|  | IB64 |  | 2012 | 0.095 | 0.082 | 0.089 |
|  | IB65 |  | 2012 | 0.051 | 0.062 | 0.057 |
|  | IB66 |  | 2012 | 0.065 | 0.066 | 0.066 |
|  | IB74 |  | 2012 | 0.062 | 0.063 | 0.063 |
|  | IB75 |  | 2012 | 0.113 | 0.112 | 0.113 |
|  | IB77 |  | 2012 | 0.063 | 0.062 | 0.063 |
|  | IB78 |  | 2012 | 0.132 | 0.146 | 0.139 |
|  | IB79 |  | 2012 | 0.137 | 0.121 | 0.129 |
|  | IB80 |  | 2012 | 0.108 | 0.103 | 0.106 |
|  | IB81 |  | 2012 | 0.089 | 0.093 | 0.091 |
|  | IB83 |  | 2012 | 0.121 | 0.118 | 0.120 |
|  | IB84 |  | 2012 | 0.106 | 0.102 | 0.104 |
|  | IB85 |  | 2012 | 0.059 | 0.055 | 0.057 |
|  | IB86 |  | 2012 | 0.066 | 0.067 | 0.067 |
|  | IB89 |  | 2012 | 0.064 | 0.062 | 0.063 |
|  | IB90 |  | 2012 | 0.100 | 0.139 | 0.120 |
|  | IB22 |  | 2012 | 0.066 | 0.063 | 0.065 |
|  | IB23 |  | 2012 | 0.068 | 0.068 | 0.068 |
|  | IB35 |  | 2012 | 0.063 | 0.061 | 0.062 |
|  | IB69 |  | 2012 | 0.069 | 0.068 | 0.069 |
|  | IB82 |  | 2012 | 0.059 | 0.060 | 0.060 |
| **CDC\*** | CDC233 | **DENV+** | 4/28/20 | 0.013 | 0.013 | 0.013 |
|  | CDC255 | **DENV+** | 4/28/20 | 0.010 | 0.009 | 0.010 |
|  | CDC269 | **DENV+** | 4/28/20 | 0.014 | 0.006 | 0.010 |
|  | CDC713 | **DENV+** | 4/28/20 | 0.052 | 0.050 | 0.051 |
|  | CDC736 | **DENV+** | 4/28/20 | 0.082 | 0.080 | 0.081 |
|  | CDC315 | **ZIKV +** | 4/28/20 | 0.058 | 0.052 | 0.055 |
|  | CDC324 | **ZIKV +** | 4/28/20 | 0.017 | 0.024 | 0.021 |
|  | CDC432 | **ZIKV +** | 4/28/20 | 0.011 | 0.012 | 0.011 |
|  | CDC493 | **ZIKV +** | 4/28/20 | 0.075 | 0.064 | 0.069 |
|  | CDC518 | **ZIKV +** | 4/28/20 | 0.055 | 0.062 | 0.059 |
|  | CDC101 | **Influenza A+** | 4/28/20 | 0.011 | 0.018 | 0.014 |
|  | CDC102 | **Influenza A+** | 4/28/20 | 0.023 | 0.023 | 0.023 |
|  | CDC103 | **Influenza A+** | 4/28/20 | 0.042 | 0.045 | 0.044 |
|  | CDC104 | **Influenza A+** | 4/28/20 | 0.015 | 0.020 | 0.017 |
|  | CDC105 | **Influenza B+** | 4/28/20 | 0.064 | 0.064 | 0.064 |
|  | CDC106 | **Influenza B+** | 4/28/20 | 0.257 | 0.255 | 0.256 |
|  | CDC107 | **Influenza B+** | 4/28/20 | 0.077 | 0.137 | 0.107 |
|  | CDC108 | Influenza B+ | 4/28/20 | 0.035 | 0.036 | 0.035 |
|  | CDC109 | Influenza B+ | 4/28/20 | 0.014 | 0.017 | 0.015 |
|  | CDC110 | Influenza B+ | 4/28/20 | 0.080 | 0.022 | 0.051 |
|  | CDC111 | Influenza B+ | 4/28/20 | 0.093 | 0.101 | 0.097 |
|  | CDC112 | Influenza B+ | 4/28/20 | 0.058 | 0.063 | 0.060 |
|  | CDC113 | RSV | 4/28/20 | 0.058 | 0.070 | 0.064 |
|  | CDC114 | RSV | 4/28/20 | 0.036 | 0.034 | 0.035 |
|  | CDC115 | RSV | 4/28/20 | 0.050 | 0.050 | 0.050 |
|  | CDC116 | RSV | 4/28/20 | 0.022 | 0.025 | 0.024 |
|  | CDC117 | RSV | 4/28/20 | 0.098 | 0.101 | 0.100 |
|  | CDC118 | RSV | 4/28/20 | 0.036 | 0.037 | 0.037 |
| **Virology**<br>**Serum Bank** | 1 |  | 4/26/17 | 0.061 | 0.069 | 0.065 |
|  | 3 | FluA H1N1+ | 6/26/18 | 0.039 | 0.041 | 0.040 |
|  | 5 | DENV + | 10/11/17 | 0.092 | 0.078 | 0.085 |
|  | 7 | DENV + | 8/17/14 | 0.024 | 0.029 | 0.026 |
|  | 9 | DENV + | 6/24/16 | 0.060 | 0.062 | 0.061 |
|  | 29 | ZIKV + | 6/29/16 | 0.027 | 0.022 | 0.024 |
|  | VB83 | ZIKV + | 2016 | 0.014 | 0.009 | 0.012 |
|  | VB84 | ZIKV + | 2016 | 0.012 | 0.016 | 0.014 |
| **Mycoplasma**<br>**IgGM+** | 107 | **Mycoplasma IgGM+** | 4/28/20 | 0.0289 | 0.0309 | 0.0299 |
|  | 123 |  | 4/28/20 | 0.0472 | 0.0456 | 0.0464 |
|  | 124 |  | 4/28/20 | 0.0313 | 0.0331 | 0.0322 |
|  | 125 |  | 4/28/20 | 0.1966 | 0.1743 | 0.1854 |
|  | 126 |  | 4/28/20 | 0.0499 | 0.0554 | 0.0526 |
|  | 127 |  | 4/28/20 | 0.0641 | 0.0804 | 0.0722 |
|  | 128 |  | 4/28/20 | 0.0516 | 0.0536 | 0.0526 |
|  | 129 |  | 4/28/20 | 0.0286 | 0.0402 | 0.0344 |
|  | 130 |  | 4/28/20 | 0.0323 | 0.0315 | 0.0319 |
|  | 131 |  | 4/28/20 | 0.0631 | 0.0738 | 0.0684 |
| **Virology**<br>**Serum Bank** | 119 | **Chikungunya +** | 5/4/20 | 0.0067 | 0.0445 | 0.0256 |
|  | 20 |  | 4/15/20 | 0.1159 | 0.0982 | 0.1070 |
|  | 118 |  | 5/4/20 | 0.0184 | 0.0281 | 0.0232 |

\* All these samples were collected prior to December 2019 in Puerto Rico. The data showed is the date in the panel of those samples was assembled at CDC Dengue Branch in support this study.

**Supplementary Table-3.**
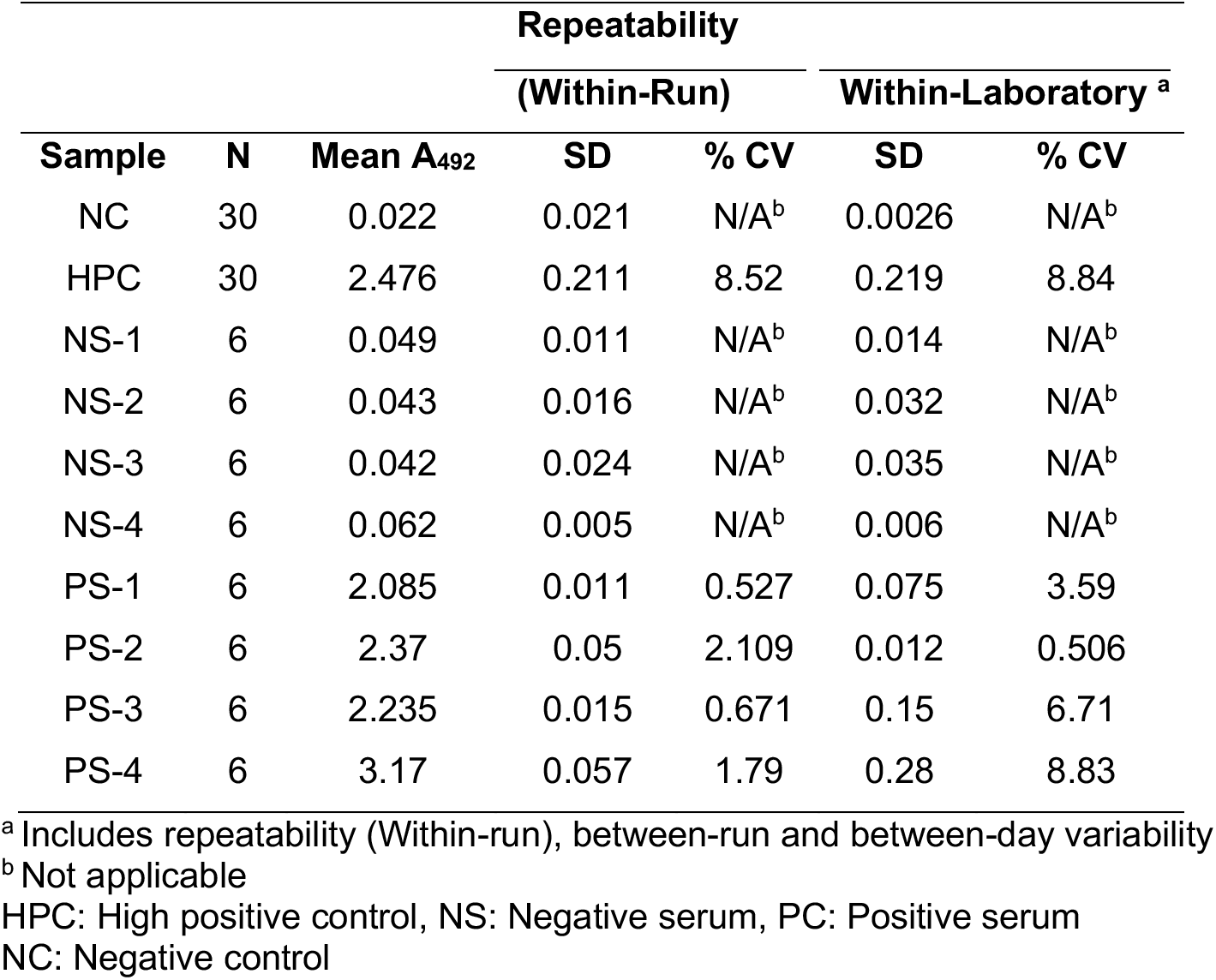
Reproducibility of CovIgG-Assay.

**Supplementary Table-4.**
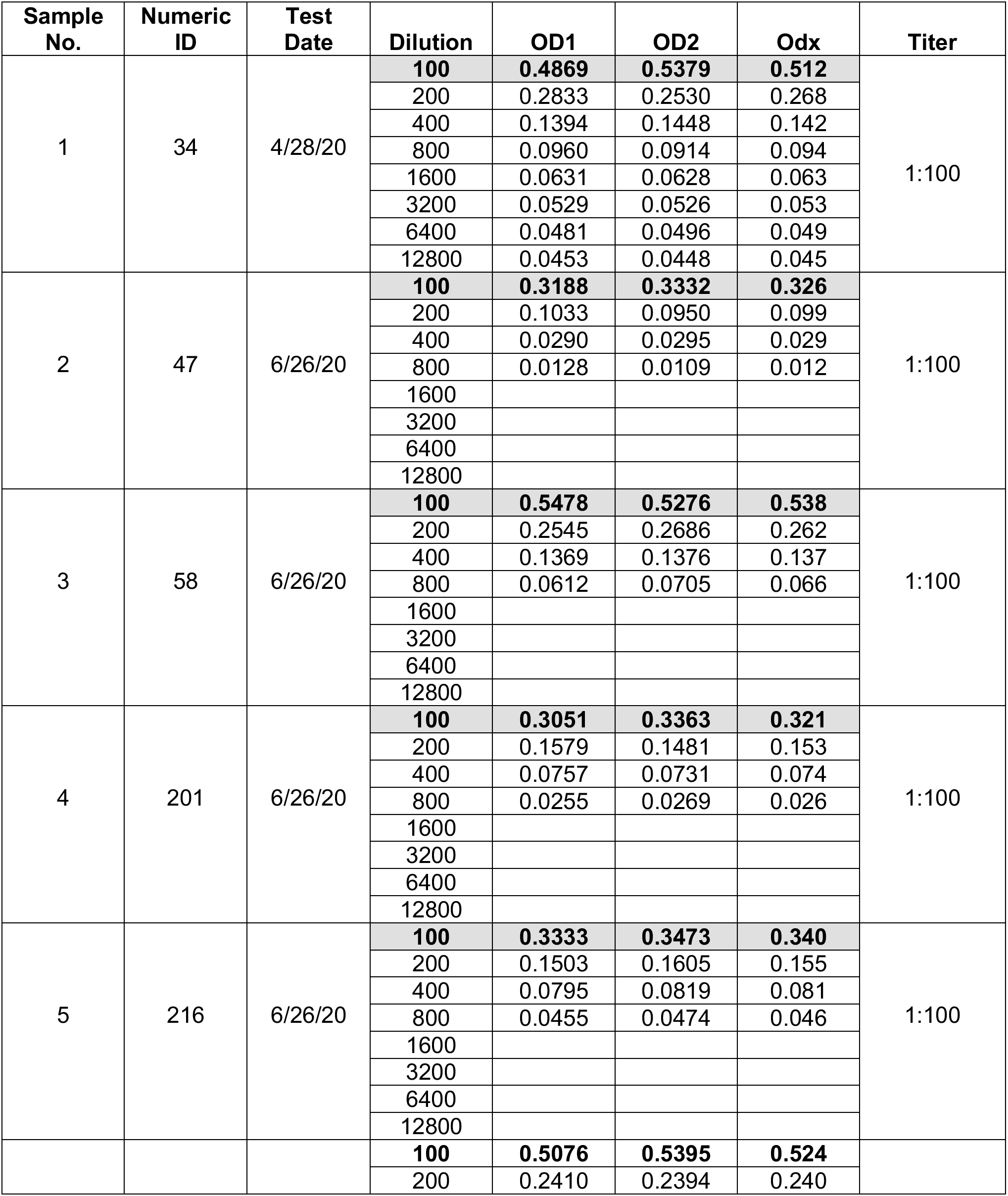
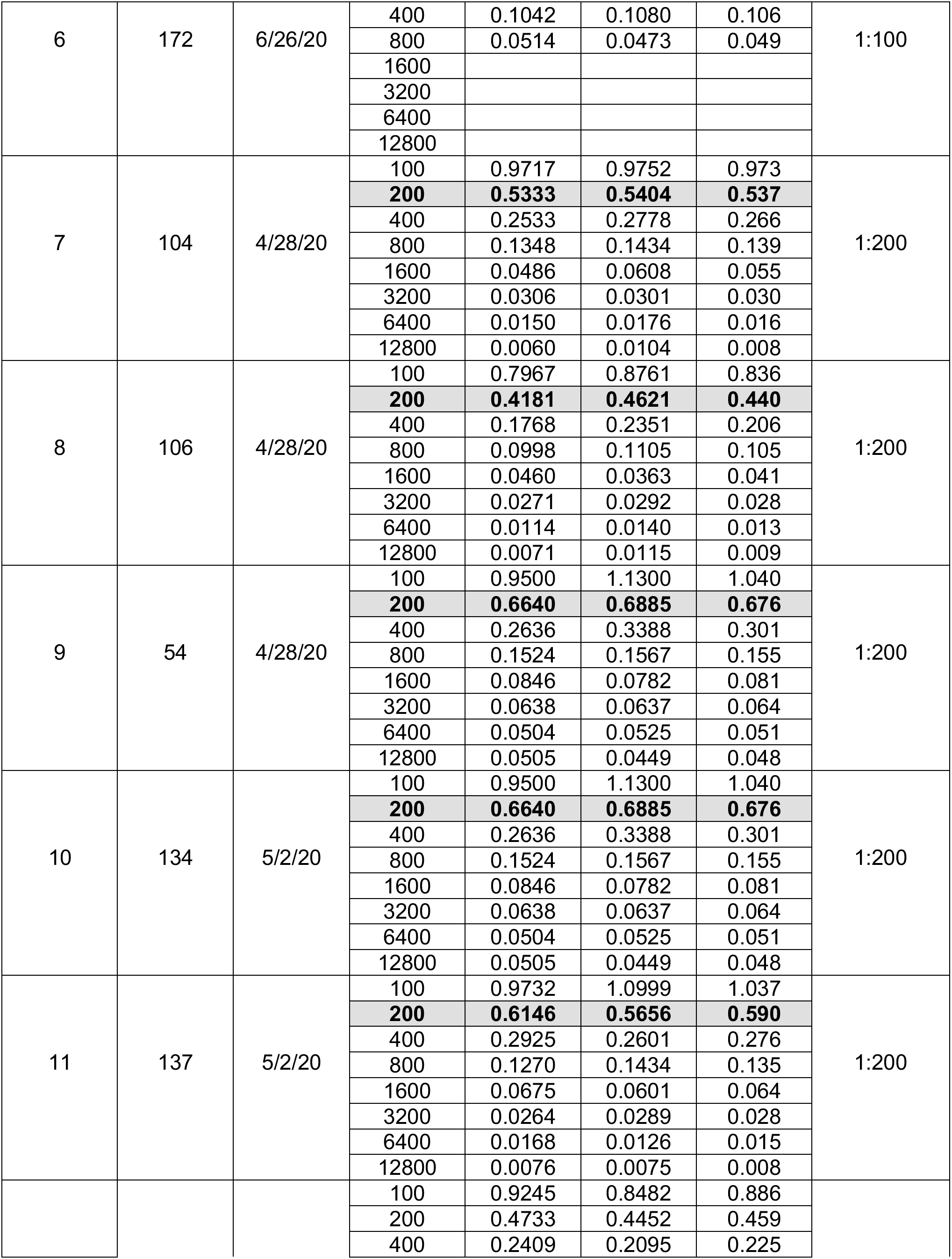
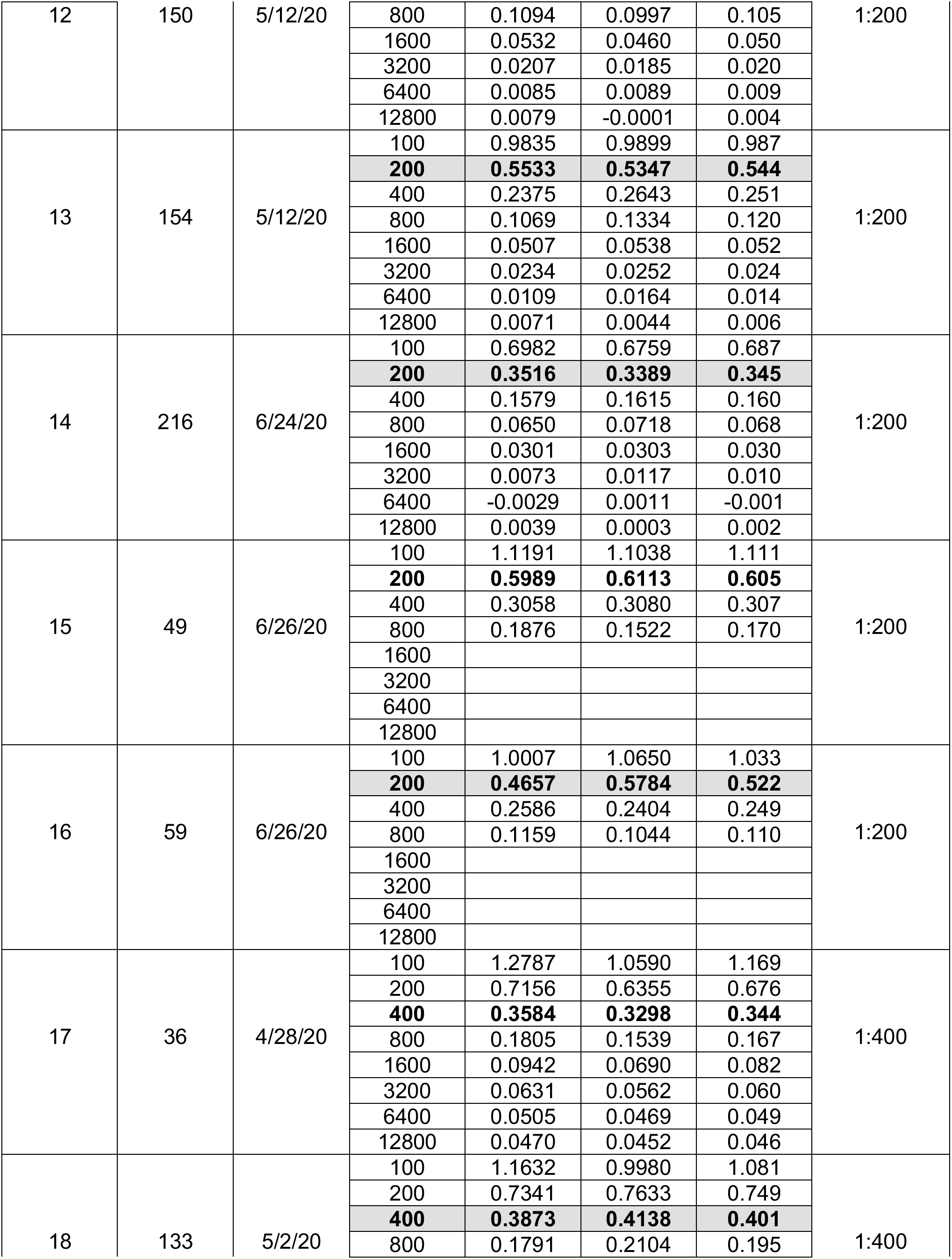
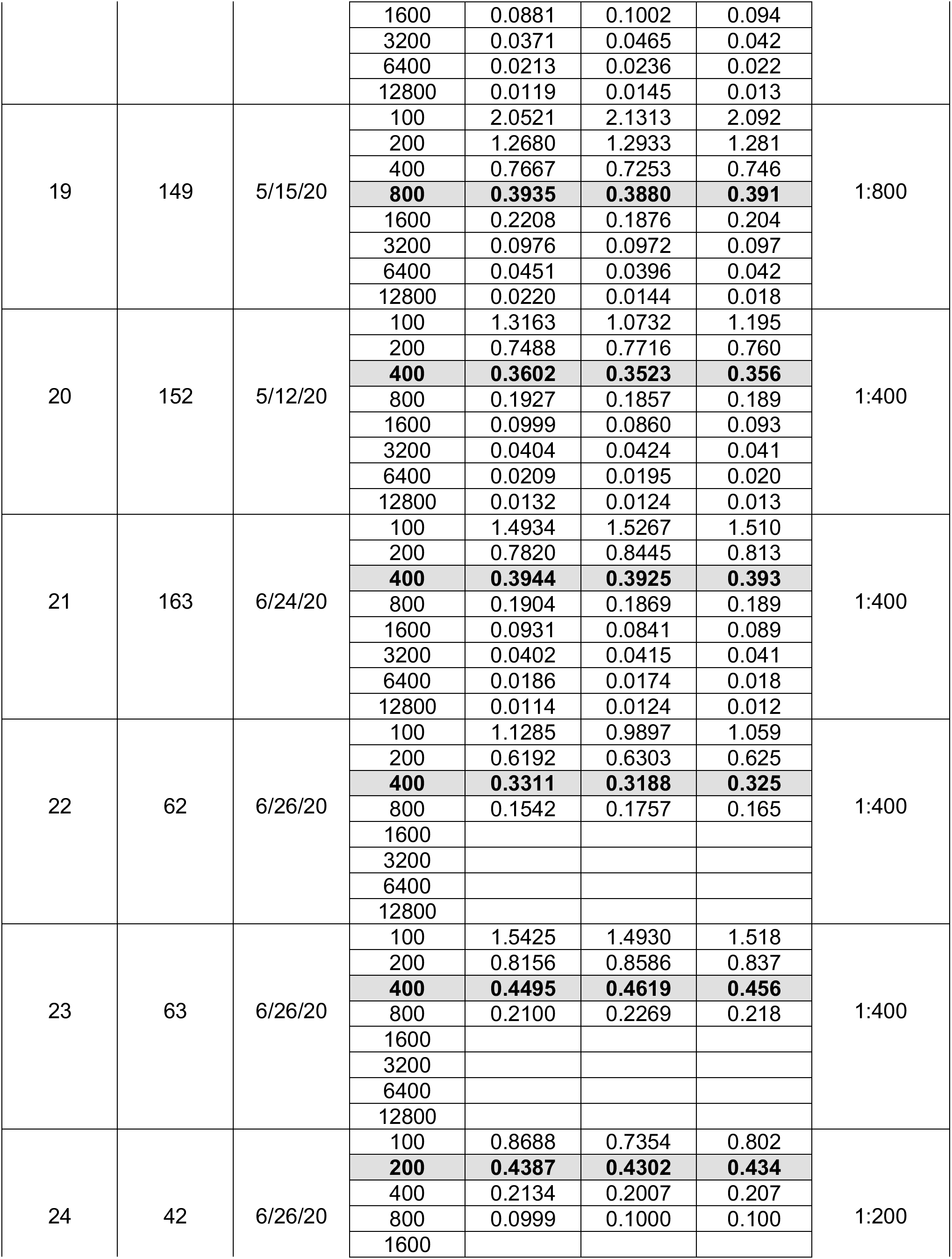
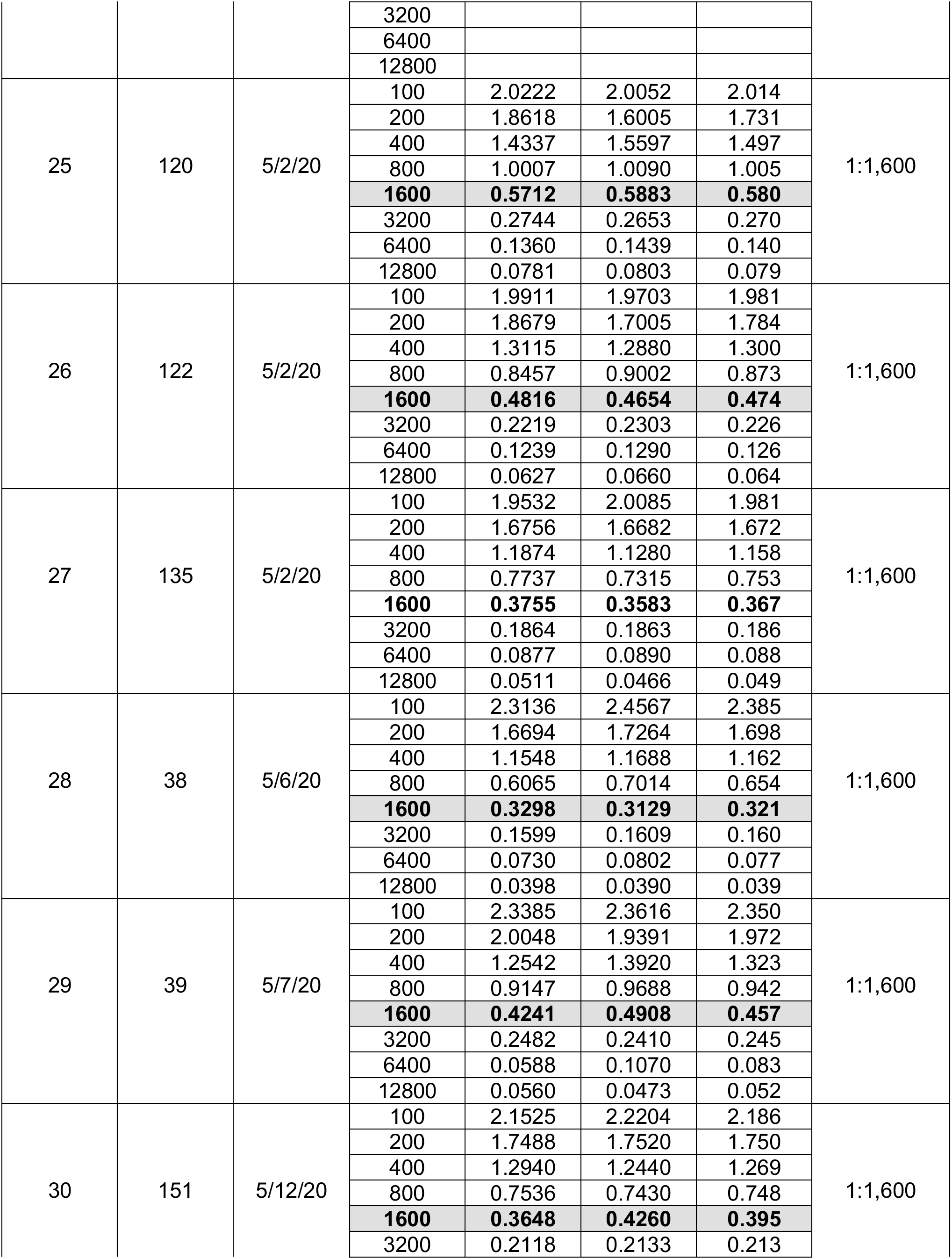
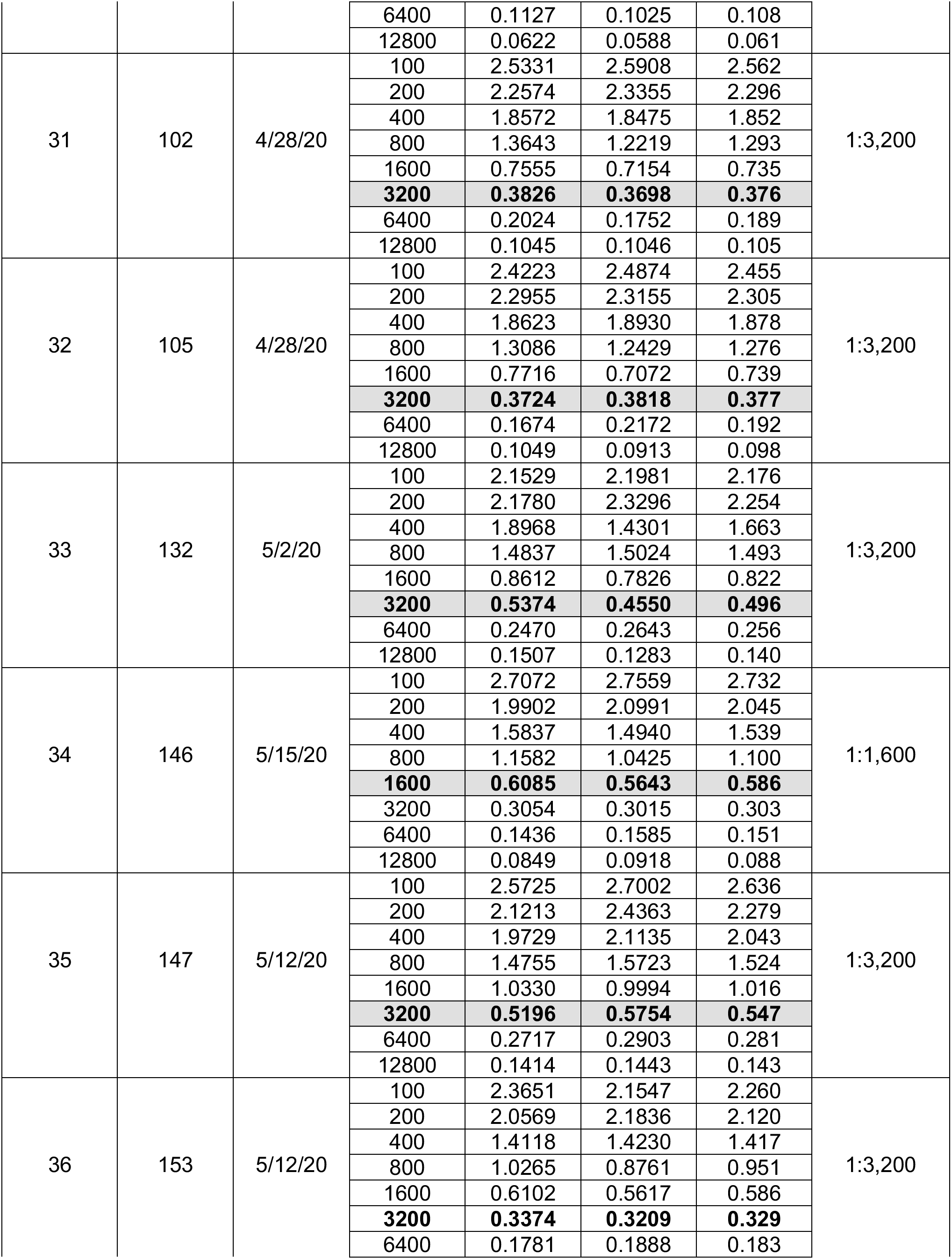
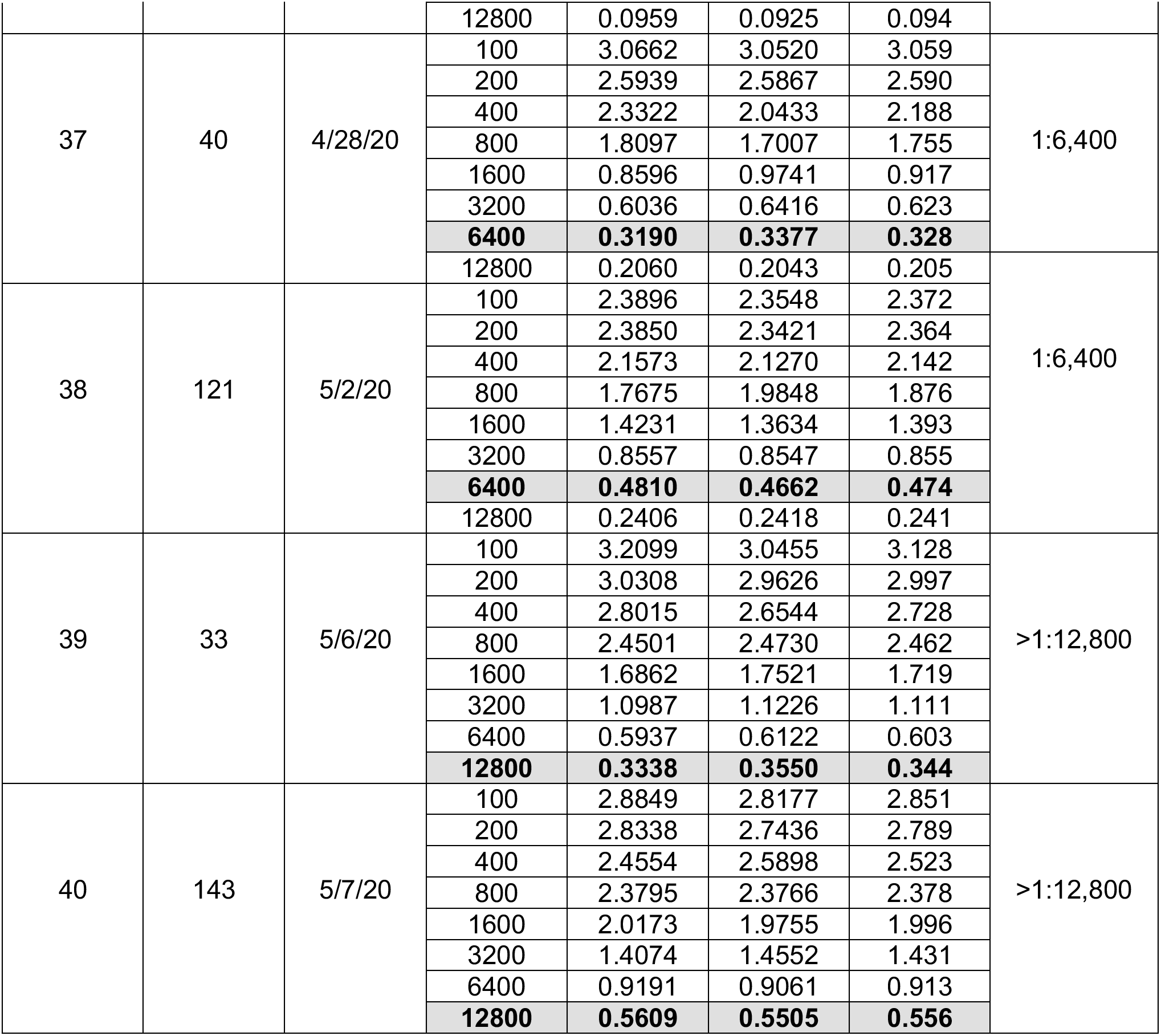
Samples titrated for establishing lineal correlation between OD (1:100) and antibody titer.

## Notes

https://prsciencetrust.org/the-covigg-assay-kit/

### Competing Interest Statement

The authors have declared no competing interest.

### Summary of Updates

Quantification of IgG was improved by end-point dilution of additional samples. Also, a correlation with another FDA EUA serological assay was added and discussed. An extended Table 5, showing the comparison of three serological tests against CoVIgG-Assay for the detection of anti-SARS CoV-2 antibodies was added. Figure 3 was corrected by adding the results of testing samples from subjects confirmed to be exposed to the Respiratory Syncytial Virus.

